# MOSAIC: A Spectral Framework for Integrative Phenotypic Characterization Using Population-Level Single-Cell Multi-Omics

**DOI:** 10.64898/2026.02.10.705077

**Authors:** Chang Lu, Yuval Kluger, Rong Ma

## Abstract

Population-scale single-cell multi-omics offers unprecedented opportunities to link molecular variation to human health and disease. However, existing methods for single-cell multi-omics analysis are either cell-centric, prioritizing batch-corrected cell embeddings that neglect feature relationships, or feature-centric, imposing global feature representations that overlook inter-sample heterogeneity. To address these limitations, we present MOSAIC, a spectral framework that learns a high-resolution feature *×* sample joint embedding from population-scale single-cell multi-omics data. For each individual, MOSAIC constructs a sample-specific coupling matrix capturing complete intra- and cross-modality feature interactions, then projects these into a shared latent space via spectral decomposition. The joint feature *×* sample embedding defines each feature’s connectivity profile per sample, enabling three downstream applications. Differential Connectivity analysis identifies features with regulatory network rewiring across conditions even when their abundance remains unchanged, revealing rewiring of proliferation programs in activated T cells from a vaccination cohort. Unsupervised subgroup detection isolates coherent feature modules to discover hidden patient subtypes, uncovering a stress-driven neuronal subtype within an HIV+ cohort. Clinical outcome prediction using connectivity-derived features complements abundance-based analysis, improving COVID-19 severity classification when integrated. MOSAIC provides a general-purpose framework for systems-level phenotypic characterization, bridging network-level discovery with clinical outcome prediction in population-scale single-cell studies.

## 1 Introduction

The proliferation of single-cell technologies has created a new frontier in biological research, with datasets of unprecedented complexity. Modern single-cell studies are expanding along two critical axes. The first is multi-modality: we can now simultaneously profile the transcriptome, epigenome, and proteome from the same cell, capturing a multi-layered view of cell states [1–7]. The second is population scale: studies now span dozens or even hundreds of individuals from large clinical cohorts and atlases, aiming to disentangle disease- or condition-driven heterogeneity from technical noise [8–12]. This dual complexity, both multi-modal and multi-sample, is essential for linking cellular phenotypes to their underlying drivers, whether disease states, genetic variation, or environmental exposures. Yet realizing this potential requires computational frameworks that can integrate diverse data modalities while explicitly accounting for the biological variation across individuals.

Current integrative frameworks can be broadly grouped by their primary analytical focus. The majority are cell-centric: their principal goal is to learn a unified, low-dimensional representation of all cells [13–21]. To do this, they harmonize information from all modalities while simultaneously correcting for sample-specific batch effects. These methods are powerful for tasks such as cell type annotation and trajectory inference across samples. However, by treating cells as the basic unit of integration, they use features as passive inputs and fail to explicitly model feature-level heterogeneity or capture how feature relationships vary across biological conditions. Conversely, feature-centric frameworks have been developed to explicitly model the rich structure of cross-modal feature interactions [22–25]. Within this domain, feature co-embedding strategies are particularly prevalent, utilizing shared latent spaces to link diverse modalities [26–28, 18, 29]. While these approaches move beyond a single-modality view, they typically generate a fixed, global feature embedding shared by all cells and all samples. This ‘one-size-fits-all’ representation, however, fundamentally masks the patient-to-patient heterogeneity that is essential for clinical insight. Thus, a critical gap remains for a framework that is both feature-centric and sample-aware: one that models how each feature’s regulatory context varies across individuals, and translates these differences into concrete biological and clinical insights, from detecting network rewiring events to stratifying patients by disease mechanism.

Here, we present MOSAIC (Multi-Omic Sample-wise Analysis of Inter-feature Connectivity), a novel spectral framework designed to fill this critical gap by learning a high-resolution feature × sample joint embedding. MOSAIC’s process is two-fold. First, it constructs a distinct joint multi-modality feature embedding for each individual in the cohort, built by modeling the intricate web of feature-feature relationships that defines that sample’s unique network structure. Second, it projects all of these individual-specific embeddings into a common latent space. The result is a powerful new data representation where a feature’s “state” is defined by its position relative to all other features, which captures its complete functional context. Because all samples are projected into a shared space, the network topology of each feature can be directly compared across samples. This embedding structure provides a unified foundation for three downstream applications: differential connectivity analysis, unsupervised sample subgroup detection, and clinical outcome prediction.

First, MOSAIC is designed to move beyond conventional differential feature detection. While powerful, existing methods [30–33] primarily rely on marginal tests, defining a “differential feature” as one whose mean level (expression, accessibility, or abundance) changes between conditions. This framework overlooks a subtler mode of variation: the rewiring of regulatory networks. MOSAIC is uniquely suited to detect such events. By defining a feature’s “state” through its functional context, MOSAIC enables Differential Connectivity (DC) analysis which tests for statistically significant shifts in a feature’s embedding position between conditions. This allows us to identify features whose functional roles and regulatory relationships are altered between conditions even when their abundance remains stable.

Second, MOSAIC can be used to identify hidden patient subgroups. In clinical studies, patients often receive a single label based on one measurement, like ‘HIV+’ or ‘diabetic’ [12, 34]. But this label can obscure important biological differences, raising the question of whether biologically meaningful subgroups exist within these seemingly uniform populations. Existing approaches [35–38] typically compute global patient similarity across all features and then cluster patients accordingly. But this ‘global’ approach often fails because critical signals from small, disease-relevant pathways may be diluted by noise from thousands of irrelevant features. MOSAIC overcomes this limitation through its feature × sample embedding. Instead of using all features simultaneously, MOSAIC first groups multi-modal features into biologically coherent modules based on their cross-patient interactions. It then tests whether any module delineates distinct patient subgroups. By isolating and evaluating these modules individually, MOSAIC reveals biologically meaningful patient subtypes that existing global methods often overlook.

Third, MOSAIC provides evidence that regulatory topology constitutes a dimension of disease variation complementary to conventional abundance measurements. MOSAIC captures the connectivity profile of each feature, encoding its relationships with all other features within each sample. Together, these per-sample connectivity profiles define a regulatory topology that can be compared across patients and used for clinical prediction. Standard approaches rely on aggregated expression levels, which capture changes in how much a gene is expressed but not in how genes interact. When connectivity-based and abundance-based features are integrated, the two sources of information are synergistic, improving predictive accuracy beyond what either achieves independently.

Through systematic benchmarking on both simulated and real multi-omic data, we validate that MOSAIC’s joint spectral integration produces biologically meaningful and cross-sample comparable embeddings. We then demonstrate three downstream applications: differential connectivity analysis, which reveals coordinated network rewiring in activated T cells from a vaccination cohort [13]; unsupervised subgroup detection, which uncovers a stress-driven neuronal subtype within HIV+ individuals [12]; and clinical outcome prediction, where MOSAIC-derived connectivity features improve COVID-19 severity classification beyond abundance-based analysis alone [39]. Together, these results establish MOSAIC as a general framework for systems-level phenotypic characterization from population-scale single-cell data.

## 2 Results

### 2.1 Overview of MOSAIC Framework

Conceptually, our framework (Figure 1A) operates in three stages. First, for each individual sample, we construct a sample-specific coupling matrix (*U*_*i*_) that captures the complete network of intra-modality and cross-modality feature-feature relationships unique to that individual. Second, we integrate these individual-specific matrices through a spectral decomposition to identify a common set of latent factors (*V*) shared across the population. Third, we project each sample’s data onto these latent factors to generate the final joint embedding of features × samples (Methods).

**Figure 1.**
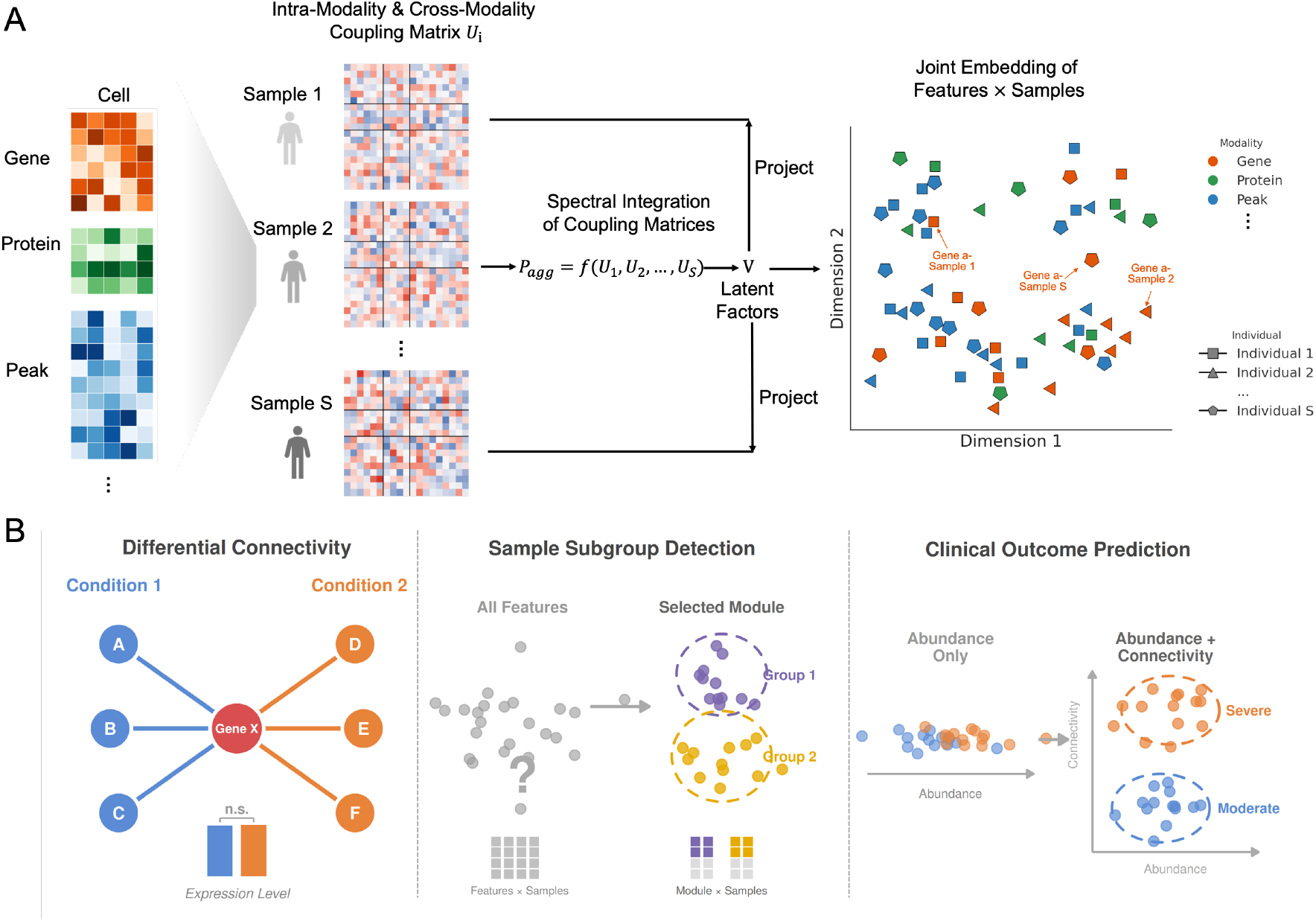
MOSAIC framework. **(A)** Schematic of the spectral integration approach: sample-specific coupling matrices (*Ui*) are constructed from multi-omic data and integrated to learn latent factors (*V*), projecting features into a shared joint embedding space. **(B)** Three downstream applications enabled by the joint embedding. *Left:* Differential Connectivity (DC) analysis identifies features whose network partners are rewired across conditions despite unchanged expression levels. *Middle:* Unsupervised sample subgroup detection isolates coherent multi-modal feature modules to reveal patient subtypes obscured by global similarity measures. *Right:* Clinical outcome prediction leverages connectivity profiles as a complementary signal to abundance, separating patient groups that expression alone cannot distinguish.

The result is a novel data representation (Figure 1A, right) in which every feature (gene, chromatin peak, or protein) is represented by an embedding vector for each sample. This vector, defined relative to all other features in the space, captures the feature’s connectivity profile within that sample. Importantly, the same feature has different embedding vectors across samples. For example, in the embedding space, Gene A has one embedding vector in Sample 1 and different vectors in Sample 2 and Sample S, reflecting how its network relationships vary between individuals. This embedding structure provides a unified foundation for three downstream applications: Differential Connectivity analysis, unsupervised sample subgroup detection, and clinical outcome prediction (Figure 1B).

### 2.2 MOSAIC Joint Embedding is Biologically Motivated and Technically Superior

A defining feature of MOSAIC is its construction of sample-specific feature embeddings within a shared latent space. To rigorously validate this framework, we structured our benchmarks around three questions (Figure 2). First, does MOSAIC’s spectral approach produce high-quality feature embeddings at a basic level (Figure 2A)? Second, is there genuine biological need for sample-specific embeddings, i.e., do feature-feature relationships truly vary across individuals (Figure 2B)? Third, given this need, does MOSAIC’s joint integration strategy produce more comparable cross-sample embeddings than the alternative of independently embedding each sample and aligning post hoc (Figure 2C, D)?

**Figure 2.**
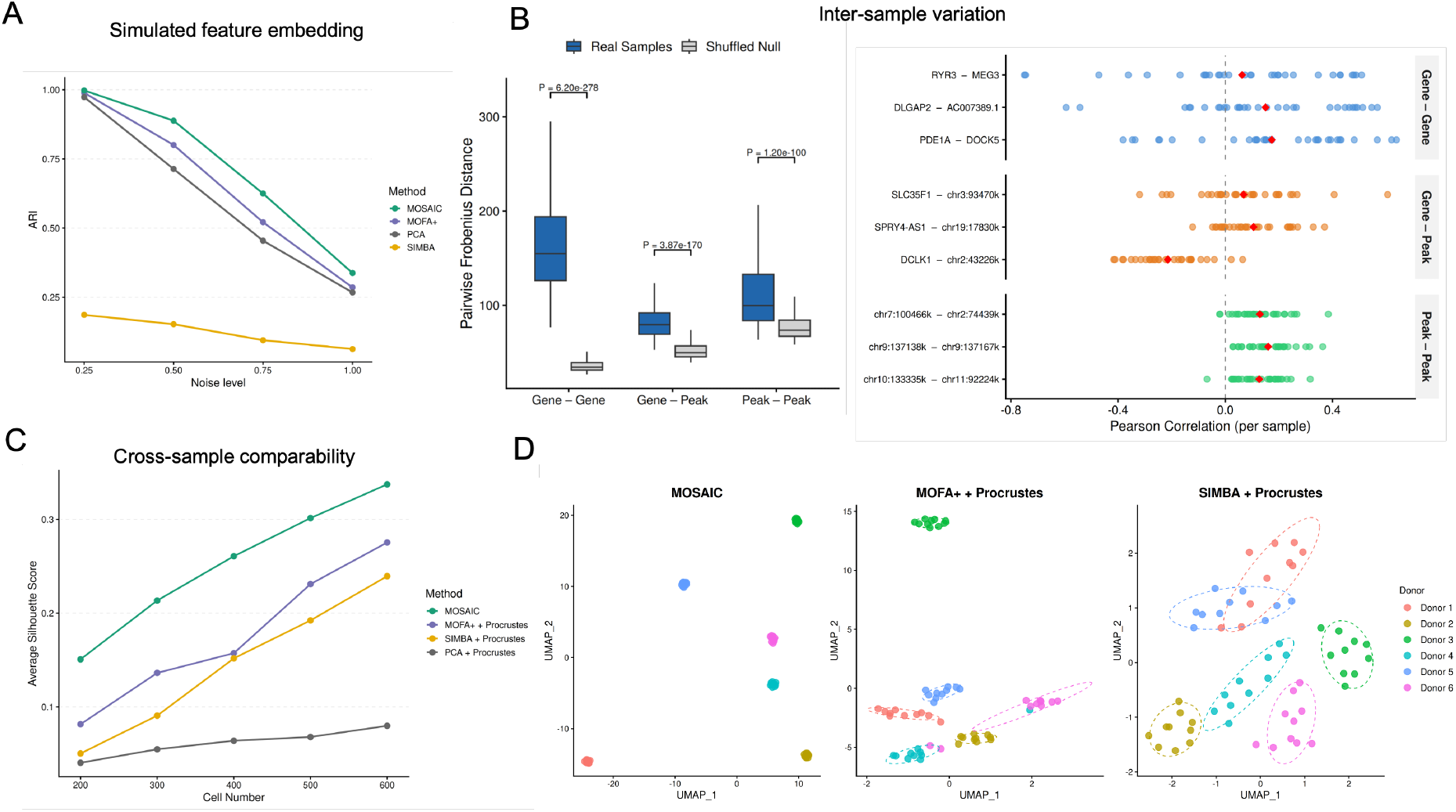
Benchmarking and biological justification of MOSAIC feature *×* sample embeddings. **(A)** ARI recovery of ground-truth feature modules on simulated multi-modal data with increasing noise levels, comparing MOSAIC, PCA, MOFA+, and SIMBA. **(B)** Left: pairwise Frobenius distances between sample-specific feature correlation matrices for real samples (blue) versus a shuffled null model (gray), stratified by modality type (gene-gene, gene-peak, peak-peak). Right: per-sample Pearson correlations for nine representative feature pairs with high cross-sample variability, grouped by modality type (red diamond: mean). **(C)** Average Silhouette Score of donor identity recovery as a function of cells per pseudo-replicate (6 healthy donors, 10 replicates each), comparing MOSAIC to MOFA+, SIMBA, and PCA each with Procrustes alignment. **(D)**UMAP visualization of pairwise embedding distances at 200 cells per replicate for MOFA+ + Procrustes, MOSAIC, and SIMBA + Procrustes. Each point represents one pseudo-replicate, colored by donor of origin (6 donors, 10 replicates each).

#### Benchmarking Feature Embedding Quality on Simulated Data

To address the first question, we assessed whether MOSAIC’s spectral approach produces high-quality feature embeddings. Because MOSAIC generates a feature *×* sample embedding tensor, we derived a single consensus feature embedding by averaging across all samples, and compared this against the global feature embeddings produced by PCA, MOFA+ [28], and SIMBA [26]. Using our multi-modal simulation framework (Supplementary Methods), we generated data with known ground-truth feature module structure and progressively increased noise levels, evaluating each method’s ability to recover the true feature groupings via the Adjusted Rand Index (ARI). At low noise levels, MOSAIC, PCA, and MOFA+ all achieved near-perfect recovery (ARI *≈* 1.0), while SIMBA showed comparatively lower performance (Figure 2A). As noise increased, all methods degraded, but MOSAIC maintained competitive performance throughout. This confirms that MOSAIC’s spectral approach is at least as effective as established methods for feature embedding, providing a solid foundation for the sample-specific evaluations that follow.

#### Feature Relationships Vary Meaningfully Across Individuals

Having established that MOSAIC produces competent feature embeddings, we next asked whether sample-specific embeddings are biologically necessary. If feature-feature relationships were largely conserved across individuals, a single global embedding would suffice and the added complexity of MOSAIC’s per-sample approach would be unwarranted. To test this, we used the multi-omic prefrontal cortex (PFC) cohort (31 donors, paired snRNA-seq and snATAC-seq) [12]. For each sample, we computed a feature-feature Pearson correlation matrix from the normalized raw data, capturing the complete network of intra- and cross-modality relationships within that individual. If feature relationships vary across individuals, these correlation matrices should differ from one another; if they are conserved, the matrices should be nearly identical. We quantified this by computing pairwise Frobenius distances between all pairs of sample-specific correlation matrices, stratified by modality block: gene-gene, gene-peak, and peak-peak (Figure 2B, left). To distinguish genuine biological variation from differences arising from finite cell sampling, we compared these distances against a null distribution generated by randomly shuffling cell-to-sample assignments, which destroys biological differences between individuals while preserving the overall data structure. Across all three modality combinations, the real inter-sample distances were significantly greater than the null expectation (Wilcoxon test, P < 0.001 for all comparisons), confirming that feature-feature relationships vary across individuals beyond what can be attributed to sampling noise. To illustrate this at the level of individual features, we selected nine representative feature pairs (three per modality type) with high cross-sample variability (Figure 2B, right). The wide spread of per-sample correlations, with some pairs shifting from positive to negative between individuals, provides direct visual evidence of the phenomenon.

#### MOSAIC Joint Integration Produces Superior Cross-Sample Comparability

The results above establish that per-sample embeddings are biologically necessary. The remaining question is how to construct them. The naive strategy is to compute an independent embedding for each sample and align them post hoc using Procrustes analysis. MOSAIC instead aggregates sample-specific coupling matrices into a shared spectral basis before projecting each sample, producing embeddings that are inherently comparable without post-hoc alignment. To evaluate which strategy better preserves biological identity across samples, we selected 6 healthy donors from the PFC cohort and split each donor’s cells into 10 non-overlapping pseudo-replicates. Pseudo-replicates from the same donor share identical biology, so a good cross-sample embedding should place them close together while separating different donors. We applied MOSAIC and three independent-then-align baselines (PCA, MOFA+, and SIMBA, each followed by Procrustes alignment) and measured donor identity recovery using the average Silhouette Score. To test robustness under data sparsity, we systematically varied the number of cells per pseudo-replicate from 200 to 600 (Figure 2C). MOSAIC consistently achieved the highest Silhouette Scores across all cell counts.

UMAP visualization at 200 cells per replicate (Figure 2D) corroborates this quantitative result: MOSAIC produced tight, well-separated donor clusters, whereas MOFA+ and SIMBA with Procrustes alignment showed considerable inter-donor overlap. Principal Coordinates Analysis (PCoA) of all four methods (Figure S2) further revealed that MOSAIC concentrated 37.6% of total variance in the first two coordinates, compared to 19.5% (MOFA+), 18.3% (SIMBA), and 8.6% (PCA), indicating that MOSAIC produces a more compact embedding structure where the leading dimensions capture the majority of cross-sample variation.

### 2.3 MOSAIC Differential Connectivity Analysis

While standard differential analysis methods focus on feature’s abundance shift, MOSAIC explicitly targets Differential Connectivity—the rewiring of a feature’s network partners independent of expression changes. The feature *×* sample embedding produced by MOSAIC is perfectly suited to detect such rewiring events. In our framework, a feature’s “state” is its functional context, defined by its position relative to all other features in the embedding space. We can therefore identify these DC features by directly testing for statistically significant shifts in a feature’s embedding position between conditions. To do this, we apply a Permutational Multivariate Analysis of Variance (PERMANOVA) test on the collection of a feature’s embedding vectors from all samples (Methods).

#### MOSAIC Uniquely Detects Differential Connectivity Events

To validate the specificity of our integrative approach and benchmark its performance as a true DC detector, we designed a comprehensive multi-modal simulation framework (Supplementary Methods). We generated data with three distinct signal types to systematically test MOSAIC against established abundance-based differential methods (DESeq2 [30], edgeR [31], MAST [32], and Seurat-Wilcoxon [40]): (1) Pure Connectivity Signal (Rewiring Only): Features had their underlying connectivity profile permuted between conditions, but their mean abundance was computationally rescaled to be identical, making this signal invisible to abundance-only methods and isolating connectivity changes alone. (2) Confounded Signal (Rewiring + Mean Shift): Features were rewired, and the resulting side-effect on mean abundance was not corrected. This represents a biologically common scenario where connectivity rewiring also produces a secondary mean expression shift. (3) Mean Shift Only: A classical differential expression signal where only feature abundance was changed, with no change in connectivity.

The results in Figure 3A demonstrate MOSAIC’s unique and orthogonal capabilities. In the confounded signal scenario, where rewiring co-occurs with mean shifts, abundance-based methods achieved relatively high performance (e.g., Seurat-Wilcoxon AUPRC = 0.845), appearing to successfully detect differential signals. However, when we isolated pure connectivity changes by rescaling means to be identical, these same methods failed dramatically, with AUPRCs dropped markedly (e.g., Seurat-Wilcoxon AUPRC = 0.498). This stark contrast reveals that traditional methods detect mean-shift artifacts rather than genuine connectivity changes. Their apparent success in the confounded scenario was driven entirely by the secondary abundance signal. In contrast, MOSAIC maintained consistently high performance in both the confounded (AUPRC = 0.981) and pure connectivity (AUPRC = 0.981) scenarios, demonstrating its ability to specifically detect network rewiring independent of mean abundance changes. Critically, for the “Mean Shift Only” signal, abundance-based methods excelled (e.g., MAST AUPRC = 0.998), while MOSAIC correctly reported no significant connectivity changes (AUPRC = 0.238). This result provides essential evidence of MOSAIC’s orthogonality to traditional differential expression approaches—it is not a general-purpose abundance detector, but rather a specialized tool designed to identify network rewiring. Together, these results confirm that MOSAIC is not simply a proxy for abundance but is specifically and sensitively detecting connectivity changes by analyzing all modalities simultaneously.

**Figure 3.**
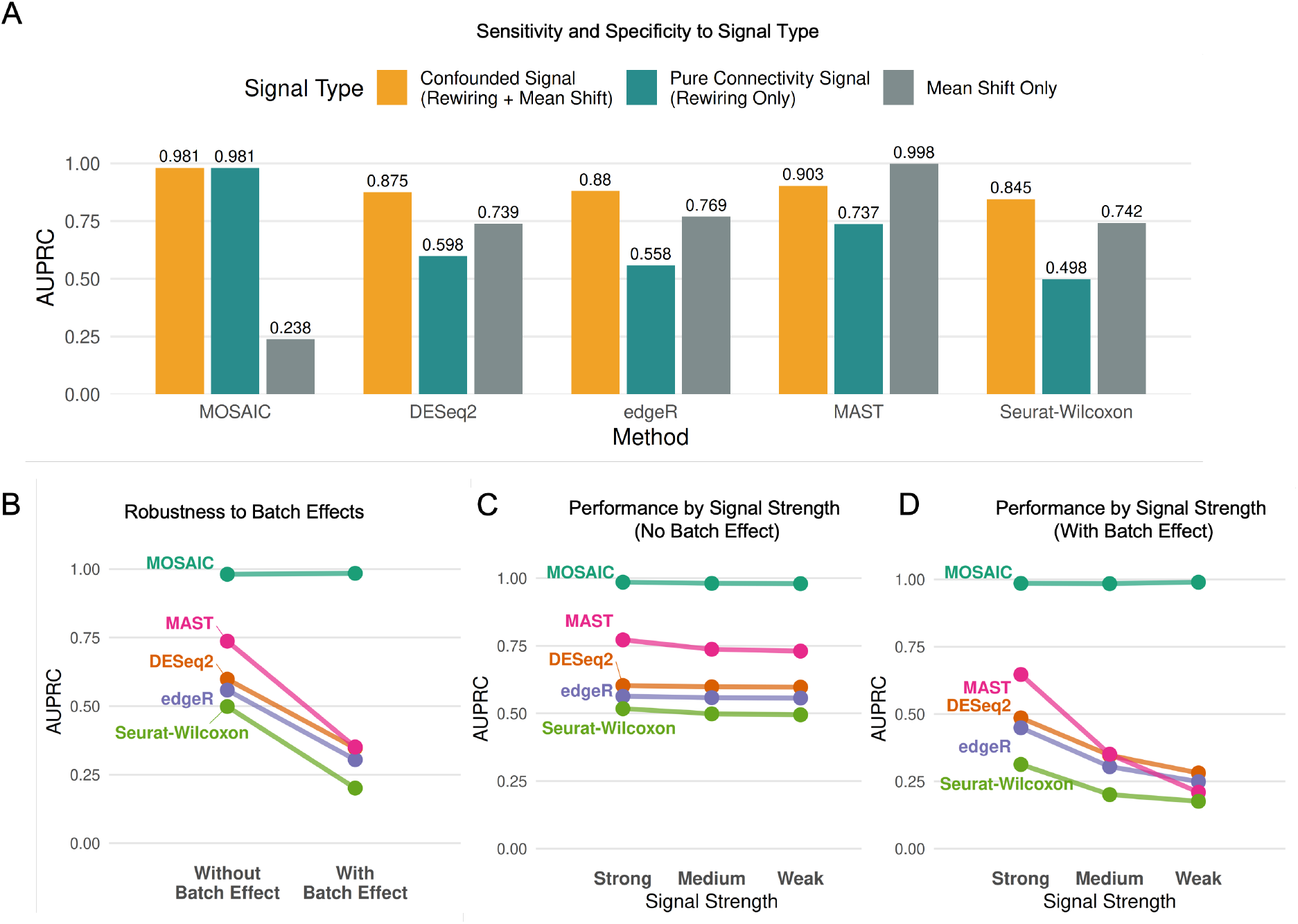
Benchmarking sensitivity and robustness. **(A)** AUPRC comparison across three simulation scenarios: Confounded Signal (Rewiring + Mean Shift), Pure Connectivity (Rewiring Only), and Mean Shift Only. MOSAIC detects pure connectivity changes missed by abundance-based methods. **(B)** Performance robustness (Slope graph) comparing methods with and without confounded batch effects. **(C–D)** Method performance with decreasing connectivity signal strength (Strong to Weak) in the absence **(C)** and presence **(D)** of batch effects.

#### MOSAIC Exhibits Robustness to Batch Effects and Signal Degradation

To demonstrate the practical utility of our approach, we evaluated its robustness against two common analytical challenges: technical batch effects and low signal-to-noise ratios.

First, we tested resilience to challenging, feature-specific batch effects in which the technical batch was confounded with the biological condition. As shown in Figure 3B, MOSAIC’s performance remained completely stable under strong batch effects (AUPRC *∼* 0.98), while all competing methods showed significant degradation (e.g., DESeq2 AUPRC dropped from 0.598 to 0.347). This demonstrates the inherent robustness of the integrative approach to confounded batch effects. Next, we assessed sensitivity using simulations with progressively weaker connectivity signals (Strong, Medium, and Weak). As signal strength decreased, MOSAIC’s accuracy remained perfect regardless of signal strength, both without (Figure 3C) and with (Figure 3D) batch effects. In contrast, competing methods revealed a critical vulnerability: while their performance was relatively stable in an idealized, no-batch-effect setting (Figure 3C), the introduction of technical noise caused severe degradation (Figure 3D). As signal strength decreased from Strong to Weak in the presence of batch effects, traditional methods approached random chance, demonstrating that their sensitivity is not robust to the combined challenge of batch effect and signal degradation, where MOSAIC still remains highly stable.

Taken together, these simulations demonstrate that MOSAIC is a robust and highly specific framework for DC analysis. It is uniquely capable of identifying pure connectivity changes that are entirely missed by traditional abundance-based methods, while remaining exceptionally resilient to the technical confounders and signal degradation that compromise other approaches.

#### MOSAIC Uncovers a Rewired Proliferation and DNA Repair Program in Activated T Cells

To demonstrate MOSAIC’s utility on real-world data, we applied it to a multi-omic CITE-seq dataset (paired measurement of transcriptomes and cell-surface proteins) of human T-cells from 8 donors[13], comparing the naive state (Day 0) to the activated state (Day 7) post-vaccination. We focused our analysis on CD4+ Naive T-cells to isolate the specific signals of T-cell activation. MOSAIC identified 393 unique DC features (p-adj < 0.05), the vast majority of which (392/393) were not found by traditional differential expression analysis by MAST. This indicates that our method uncovers a completely distinct layer of biological change. To validate that these unique hits were biologically meaningful and not random noise, we performed three complementary global analyses.

First, Reactome pathway analysis [41] revealed that the 393 DC features are significantly enriched in pathways central to T-cell clonal expansion (Figure 4A). The top enriched terms delineate a coordinated program spanning “Cell Cycle Checkpoints” and the “G1/S Transition,” alongside critical DNA repair mechanisms such as “HDR through Homologous Recombination.” This confirms that the features rewired during activation represent the core machinery linking rapid cell proliferation with the genome integrity mechanisms necessary to support it. Second, protein-protein interaction network analysis using STRING revealed that the DC proteins showed significant enrichment for known physical and functional interactions (493 interactions observed vs. 392 expected, *p* = 5.56*e* − 07; Figure S1). This network enrichment indicates that the rewired features are not randomly distributed but form an interconnected regulatory network, providing additional evidence that MOSAIC identifies functionally coordinated sets of features undergoing concerted reorganization during T-cell activation. Third, we quantitatively validated that DC features exhibit significantly higher network turnover than background features across condition. We measured the stability of each feature’s functional context by calculating the Jaccard Index of its neighbor overlap between the naive (Day 0) and activated (Day 7) states (Figure 4B). The DC features showed significantly lower Jaccard scores (*p* = 4.3 × 10^−15^), providing statistical evidence that their regulatory neighborhoods undergo substantially greater reorganization compared to the global background.”

**Figure 4.**
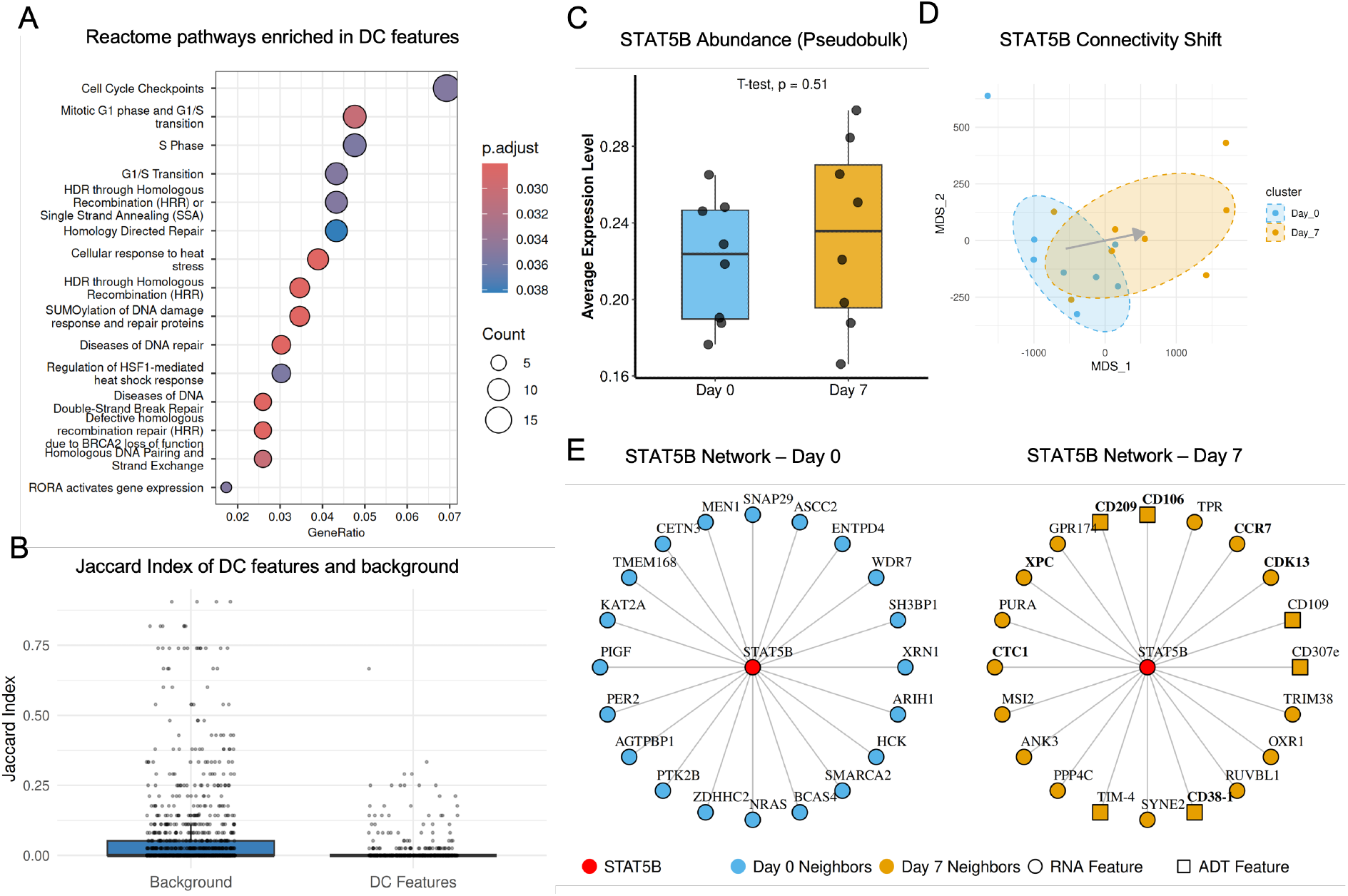
Rewiring of proliferation networks in activated T cells. **(A)** Reactome pathway enrichment of 393 DC features. **(B)** Jaccard Index of neighbor overlap (Day 0 vs. Day 7) for DC features versus background, showing higher turnover in DC features (*p* = 4.3 *×*10^*−*15^). **(C)** Box plot of pseudobulk expression (average per sample) showing no significant difference between Day 0 and Day 7 (*p* = 0.51, T-test). **(D)** MOSAIC embedding shift of STAT5B from Day 0 (blue) to Day 7 (orange), showing systematic shift in the mean connectivity profile. **(E)** STAT5B’s top 20 connectivity partners at Day 0 (left) and Day 7 (right), showing network topology turnover.

To illustrate these findings at a mechanistic level, we investigated the STAT5B, a canonical T-cell transcription factor identified by our method (DC rank 93, p-adj = 0.015). Traditional methods missed this key regulator, as its gene expression level remained stable between Day 0 and Day 7 (Figure 4C). MOSAIC, however, detected a highly significant shift in its connectivity profile, visualized by the clear separation of its sample embeddings (Figure 4D), with an arrow indicating the systematic shift from the Day 0 centroid to the Day 7 centroid. To interpret the biological meaning of this rewiring, we examined the top 20 connectivity partners of STAT5B (Figure 4E). The network exhibited a complete turnover in partners between the two timepoints, with no overlap. The Day 0 network consisted of general regulators, such as the chromatin remodeler SMARCA2 and the signaling protein NRAS. In contrast, the Day 7 network was rewired to features directly controlling the activated T-cell phenotype. We found new connections to the canonical trafficking receptor CCR7 [42] and the classic activation marker CD38-1 [43], as well as numerous other surface proteins like CD106 and CD209. Notably, this rewiring directly links STAT5B to the proliferation machinery discovered in our pathway analysis (Figure 4A). New Day 7 partners include the cell-cycle regulator CDK13 [44] and the DNA repair proteins XPC and CTC1 [45, 46]. This aligns with recent evidence that STAT5B phosphorylation is strictly required to induce the expression of cell cycle regulators necessary for maximal T cell expansion [47]. Moreover, the concurrent recruitment of DNA repair factors reflects the critical biological requirement to maintain genomic integrity during the replication stress of rapid cell division [48]. Together, these results demonstrate that STAT5B, while not changing in abundance, has completely rewired its functional context from a basal regulatory state to one that is poised to orchestrate T-cell activation, trafficking, and proliferation—a multi-omic functional transition that is invisible to traditional abundance-based methods.

### 2.4 MOSAIC Enables Unsupervised Multi-Modal Sample Subgroup Detection

We next applied MOSAIC to dissect latent patient heterogeneity. Unlike global clustering methods that can be obscured by noise from irrelevant features, MOSAIC utilizes a modular strategy to identify robust patient subgroups. Specifically, we leverage the joint embedding to calculate a stratification profile for each feature, capturing how it distinguishes samples from one another. By clustering these profiles, MOSAIC identifies feature modules, which are groups of features that co-vary similarly across the cohort (Methods). This allows us to move beyond a single global map and evaluate whether specific molecular modules drive distinct, biologically meaningful patient subgroups.

#### Validation of MOSAIC Subgroup Detection on HIV and Control Cohorts

To first validate this approach, we applied it to L2/3 inhibitory neurons from the multi-omic prefrontal cortex cohort of 30 donors (18 HIV+, 12 controls) with paired snRNA and snATAC measurements [12], treating all disease labels as unknown. We sought to test whether MOSAIC could successfully identify the subgroup structure distinguishing HIV+ samples from controls without prior knowledge. Applying the feature-clustering workflow described above, the analysis successfully identified a prominent feature module (Figure 5A). The sample embedding based on this module shows clear and significant separation between the two cohorts (Figure 5B). To validate that this module captures true biological signal rather than technical artifacts, we calculated module scores for each cell from the original single-cell data (Supplementary Methods). The distribution of module scores differed significantly between cells from HIV+ and control samples (Figure 5C), showing a clear increase in the HIV+ cohort and confirming that the identified feature module reflects genuine HIV-associated effects in this neuronal population. Together, these results validate that MOSAIC’s feature-centric approach can robustly identify primary biological structure in complex datasets without supervision.

**Figure 5.**
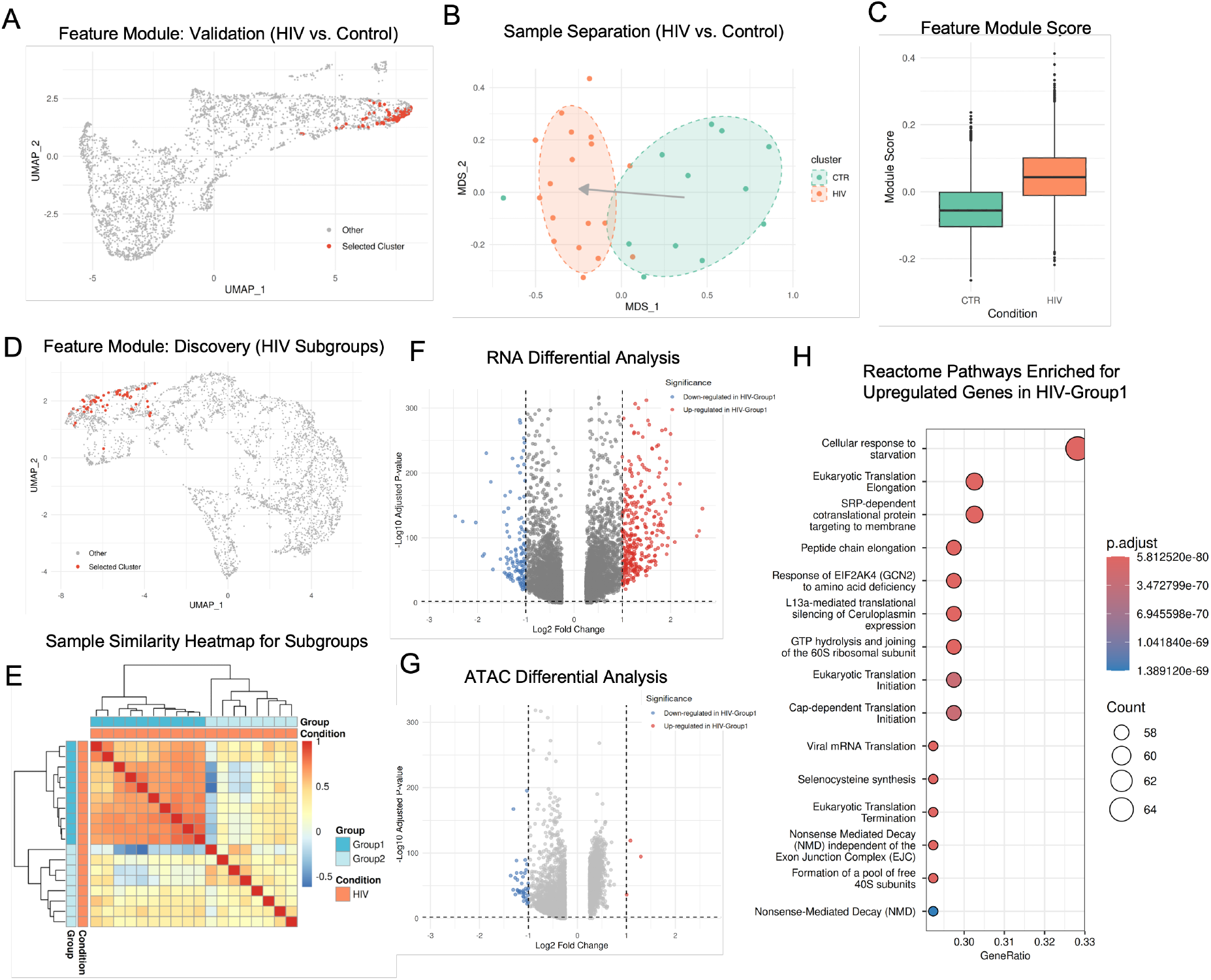
**(A–C)** Validation using L2/3 inhibitory neurons HIV+ and Control samples: **(A)** Feature module (red) identified to validate the separation of HIV+ and control samples. **(B)** MOSAIC sample embeddings based on the module in A, showing clear separation of the HIV+ and control samples. **(C)** Feature module scores in HIV+ vs. Control. **(D–H)** Discovery of HIV subgroups: **(D)** Feature module (red) identified within the HIV+ cohort. **(E)** Sample similarity heatmap based on the discovery feature module from panel D, partitioning HIV+ samples into a coherent HIV-Group1 and a heterogeneous HIV-Group2. **(F)** RNA differential expression and **(G)** ATAC differential accessibility between the two HIV subgroups. **(H)** Reactome enrichment for genes upregulated in HIV-Group 1.

#### MOSAIC Discovers a New Stress-Driven Neuronal Subtype in HIV+ Patients

The true power of this approach lies in discovering previously unknown patient subtypes hidden within diagnostic labels. To demonstrate this capability, we applied MOSAIC’s unsupervised clustering exclusively to the 18 HIV+ samples. This analysis revealed a robust feature module (Figure 5D) that partitioned the HIV+ cohort into two distinct, previously unrecognized subgroups. This cryptic structure is clearly visible in the sample similarity heatmap (Figure 5E). Based solely on this feature module, the samples were clustered into two groups: HIV-Group1 (n=10) forms a tight, highly coherent cluster, while HIV-Group2 (n=8) appears more heterogeneous. This structure suggests that the feature module has identified a common biological state shared by patients in Group 1 that is absent or variable in Group 2.

To validate this hypothesis and characterize this putative state, we performed differential analysis between the two groups. The results confirmed a strong, RNA-dominated biological shift. The RNA volcano plot (Figure 5F) shows a dramatic asymmetry, with numerous genes significantly upregulated in HIV-Group1 compared to HIV-Group2. In contrast, this massive gene-level signal was not mirrored at the chromatin level, where we observed very few significantly differential ATAC features (Figure 5G). This confirms that Group 1 is defined by the gain of a specific transcriptional program that is absent in Group 2. To define the functional identity of this state, we performed Reactome pathway enrichment analysis on the upregulated genes in HIV-Group1. The results (Figure 5H) revealed a dominant signature of translational machinery operating under metabolic stress. The profile was driven by the top term “Cellular response to starvation,” alongside a broad upregulation of core protein synthesis components (e.g., translation elongation and SRP-dependent targeting). Notably, the specific enrichment of GCN2-mediated signaling indicates activation of the Integrated Stress Response (ISR). This molecular profile implies that HIV-Group1 neurons are undergoing chronic metabolic exhaustion and proteostatic stress, a known feature of HIV-associated neurocognitive disorders (HAND) that compromises neuronal function [49, 50]. The paradoxical upregulation of translation machinery in this context may reflect a compensatory effort to maintain synaptic plasticity despite the neurotoxic environment [51].

Together, these findings demonstrate that MOSAIC identified a biologically coherent subgroup of HIV+ samples within L2/3 inhibitory neurons defined by a specific stress-response endotype. By focusing on modular connectivity rather than global similarity, MOSAIC revealed a translation-driven, ISR-like state that would likely remain invisible to standard clustering approaches.

### 2.5 MOSAIC Uncovers Complementary Topological Mechanisms of Disease Severity

To evaluate the clinical utility of MOSAIC, we assessed whether its connectivity-based embeddings could predict patient outcomes and provide information complementary to conventional gene expression analysis. We applied MOSAIC to a large-scale COVID-19 transcriptomic dataset [39], comprising scRNA-seq data from 151 patients stratified by disease severity (Moderate vs. Severe). This dataset provides the statistical power necessary for robust outcome prediction, a scale rarely achieved in paired multiomic studies. Crucially, while MOSAIC is designed for multi-omic integration, its core mathematical framework is modality-agnostic, allowing it to unlock latent regulatory information from standard single-cell transcriptomes.

Using this framework, we first performed an unbiased computational screen across all major immune lineages captured in the dataset. This analysis identified monocytes as the most potent predictors of disease severity (AUC = 0.851; Figure 6A), outperforming T, B, and NK cell populations. We therefore focused subsequent analyses on the monocyte compartment to determine whether connectivity-based features could reveal pathology missed by standard expression analysis.

**Figure 6.**
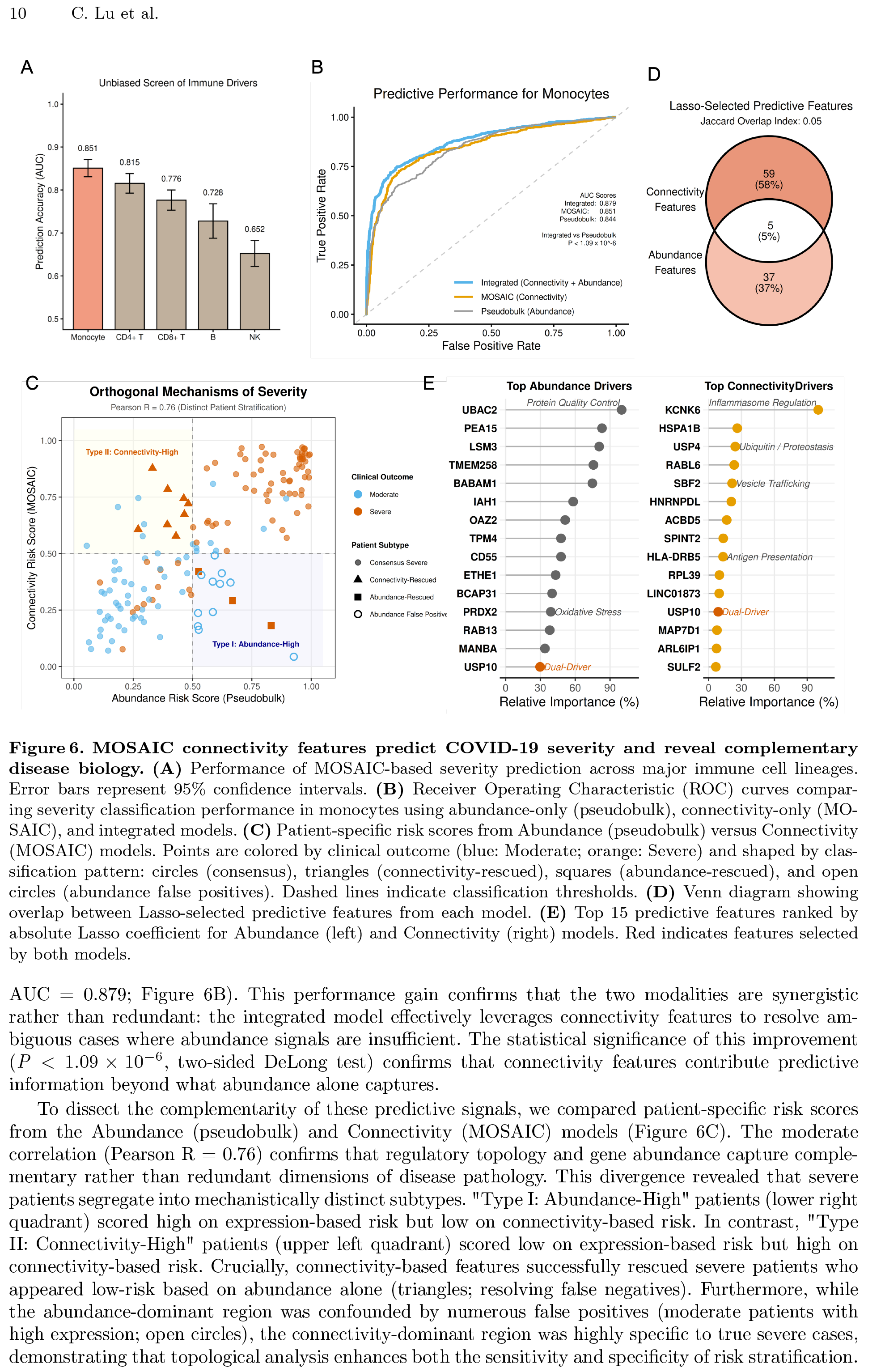
MOSAIC connectivity features predict COVID-19 severity and reveal complementary disease biology. **(A)** Performance of MOSAIC-based severity prediction across major immune cell lineages. Error bars represent 95% confidence intervals. **(B)** Receiver Operating Characteristic (ROC) curves comparing severity classification performance in monocytes using abundance-only (pseudobulk), connectivity-only (MOSAIC), and integrated models. **(C)** Patient-specific risk scores from Abundance (pseudobulk) versus Connectivity (MOSAIC) models. Points are colored by clinical outcome (blue: Moderate; orange: Severe) and shaped by classification pattern: circles (consensus), triangles (connectivity-rescued), squares (abundance-rescued), and open circles (abundance false positives). Dashed lines indicate classification thresholds. **(D)** Venn diagram showing overlap between Lasso-selected predictive features from each model. **(E)** Top 15 predictive features ranked by absolute Lasso coefficient for Abundance (left) and Connectivity (right) models. Red indicates features selected by both models.

As a baseline, we generated pseudobulk profiles by aggregating raw gene expression counts for each patient into a single feature vector, then trained a parallel Lasso classifier. While the pseudobulk baseline achieved robust performance (AUC = 0.844), MOSAIC connectivity profiling alone performed comparably (AUC = 0.851). Crucially, integrating both approaches improved predictive accuracy (Integrated AUC = 0.879; Figure 6B). This performance gain confirms that the two modalities are synergistic rather than redundant: the integrated model effectively leverages connectivity features to resolve ambiguous cases where abundance signals are insufficient. The statistical significance of this improvement (*P <* 1.09 *×* 10^*−*6^, two-sided DeLong test) confirms that connectivity features contribute predictive information beyond what abundance alone captures.

To dissect the complementarity of these predictive signals, we compared patient-specific risk scores from the Abundance (pseudobulk) and Connectivity (MOSAIC) models (Figure 6C). The moderate correlation (Pearson R = 0.76) confirms that regulatory topology and gene abundance capture complementary rather than redundant dimensions of disease pathology. This divergence revealed that severe patients segregate into mechanistically distinct subtypes. “Type I: Abundance-High” patients (lower right quadrant) scored high on expression-based risk but low on connectivity-based risk. In contrast, “Type II: Connectivity-High” patients (upper left quadrant) scored low on expression-based risk but high on connectivity-based risk. Crucially, connectivity-based features successfully rescued severe patients who appeared low-risk based on abundance alone (triangles; resolving false negatives). Furthermore, while the abundance-dominant region was confounded by numerous false positives (moderate patients with high expression; open circles), the connectivity-dominant region was highly specific to true severe cases, demonstrating that topological analysis enhances both the sensitivity and specificity of risk stratification.

We next interrogated the molecular features driving these divergent patient stratifications. We extracted the predictive genes with non-zero coefficients from the respective Lasso models and observed a striking lack of overlap (Jaccard Index = 0.05; Figure 6D), confirming that the two models rely on largely non-overlapping sets of predictive genes to reach their predictions. Functional annotation of the 15 top-weighted drivers revealed a clear mechanistic dichotomy (Figure 6E). While abundance predictors were primarily enriched for generalized oxidative stress markers (e.g., PRDX2, ETHE1, UBAC2), connectivity predictors mapped to specific immunoregulatory machinery.

Notably, the top topological driver, KCNK6 (TWIK-2), is a K^+^ efflux channel that creates the ionic environment necessary for ATP-induced NLRP3 inflammasome activation[52], linking network rewiring to the mechanism of sterile inflammation. Furthermore, MOSAIC uniquely identified genes involved in antigen presentation (HLA-DRB5) and vesicle trafficking (SBF2), processes that rely on complex protein interaction networks rather than simple stoichiometric abundance. Interestingly, the deubiquitinase USP10 emerged as a rare ‘dual-driver’ essential to both modalities, identifying proteostasis as a convergent node where cellular stress (abundance) and regulatory rewiring (connectivity) intersect.

Together, these results demonstrate that MOSAIC extracts clinically actionable network-level features from standard single-cell transcriptomes, enabling patient stratification beyond what conventional expression analysis can achieve.

## 3 Discussion

### Abundance and Connectivity Capture Complementary Dimensions of Variation

A central conceptual contribution of this work is the dissociation of a feature’s “state” (abundance) from its “function” (connectivity). Standard differential analysis identifies features by changes in expression level, but this framework overlooks regulatory rewiring, where a feature’s network context shifts despite stable abundance. Our differential connectivity analysis provides direct evidence for this dissociation at the feature level: STAT5B showed no expression change upon T-cell activation yet completely rewired its connectivity partners from basal regulators to proliferation and DNA repair machinery. The COVID-19 severity analysis extends this principle to the patient level. Abundance-based and connectivity-based risk scores showed moderate correlation (Pearson R = 0.76) with strikingly non-overlapping predictive features (Jaccard Index = 0.05), revealing that severe patients segregate into mechanistically distinct subtypes driven by either elevated expression or perturbed regulatory wiring. Together, these results establish that connectivity constitutes a complementary biological dimension, one that enhances both mechanistic discovery and clinical prediction beyond what abundance analysis alone can achieve.

### Modular vs. Global Patient Stratification

A distinct methodological contribution is MOSAIC’s modular paradigm for patient stratification. Conventional clustering, which computes global patient similarity across all features, is prone to signal dilution when disease heterogeneity is driven by small, specific pathways. By instead deriving patient similarity from coherent feature modules, MOSAIC amplifies relevant biological signals while suppressing noise from invariant features. This targeted approach is critical for decomposing complex diagnostic labels into mechanistically distinct endotypes, as demonstrated by our identification of the stress-driven “cryptic” subgroup within the HIV+ cohort.

### Robustness via Spectral Network Abstraction

MOSAIC mitigates technical noise by abstracting data into second-order feature relationships. By converting raw expression values into correlation-based coupling matrices, our framework prioritizes relative data structure over absolute magnitudes. This effectively acts as a filter for technical variation: while batch effects often manifest as global shifts in mean expression magnitudes, local feature-feature correlations remain comparatively stable. Consequently, MOSAIC achieves “zero-configuration” robustness, maintaining high specificity in the presence of confounded batch effects without requiring the explicit correction parameters often needed by abundance-based models.

### Computational Complexity and Limitations

The rigorous modeling of feature relationships imposes specific computational demands. The construction of sample-specific coupling matrices scales as *𝒪* (*S · n · F* ^2^), where *S* is the number of samples, *n* is the average number of cells, and *F* is the number of features. The subsequent spectral decomposition scales as *𝒪* (*S · F* ^2^ · *k*) by utilizing a truncated eigendecomposition for the top *k* components. Thus, the framework is computationally linear with respect to sample size (making it scalable to large cohorts) but quadratic with respect to feature count. For extremely high-dimensional spaces (eg. *F >* 50, 000), this necessitates feature selection to prioritize informative variables. Furthermore, the current framework assumes paired multi-omic data. Extending MOSAIC to unpaired modalities would require upstream manifold alignment to infer cell-cell correspondences, utilizing computational frameworks such as UnpairReg [53].

### Conclusion

MOSAIC establishes a spectral framework for analyzing the relational structure of single-cell populations. By shifting the analytical lens from abundance to connectivity, it uncovers regulatory wiring that underpins cellular states but remains invisible to standard metrics, and translates these network-level insights into clinically actionable predictions. This approach provides a technically robust, mathematically grounded toolkit for dissecting the complex phenotypic heterogeneity that defines human health and disease.

## 4 Methods

### 4.1 MOSAIC Framework

The MOSAIC framework learns a joint feature *×* sample embedding from population-level, multi-omic single-cell data through a three-stage spectral procedure: (1) constructing a sample-specific coupling matrix for each individual, (2) integrating these matrices to extract a shared latent basis, and (3) projecting each sample onto this basis to produce comparable, sample-specific feature embeddings.

#### Input Structure and Data Preprocessing

Consider a cohort of *S* samples and *K* data modalities (e.g., RNA, ATAC, Protein). For each sample *i ∈ {*1, …, *S}*, the data from modality *k* is represented by a cell-by-feature matrix 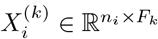, where *n*_*i*_ is the number of cells in sample *i* and *F*_*k*_ is the number of features in modality *k*. All input data are assumed to be normalized and scaled according to standard practices for each modality.

#### Construction of Sample-Specific Coupling Matrices

For each sample *i*, we concatenate all modality-specific matrices horizontally into a unified cell-by-feature matrix:

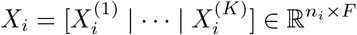

where 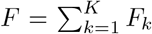 is the total number of features across all modalities. We then compute a sample-specific coupling matrix *U*_*i*_ *∈*ℝ^*F ×F*^ that encodes the complete network of intra- and cross-modality feature relationships for that individual. Each entry is defined as the cosine similarity between two feature vectors (columns of *X*_*i*_):

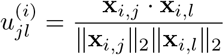

where **x**_*i,j*_ denotes the *j*-th column of *X*_*i*_. The use of cosine similarity normalizes for differences in feature magnitude across modalities, ensuring that intra-modality (e.g., gene-gene) and cross-modality (e.g., gene-peak) relationships are measured on a comparable scale. The resulting matrix *U*_*i*_ is symmetric and positive semi-definite, representing the complete inter-feature connectivity topology specific to sample *i*.

#### Spectral Integration and Latent Factor Generation

The goal of this stage is to identify latent factors shared across the population, onto which individual samples can be projected to produce comparable embeddings. A naive approach would eigendecompose each *U*_*i*_ independently and align the resulting eigenvectors post hoc (e.g., via Procrustes analysis). However, independent decomposition suffers from inherent sign and rotation ambiguity, making cross-sample alignment ill-posed. MOSAIC instead adopts an aggregate-then-decompose strategy: by pooling information across all samples before extracting latent factors, the shared spectral structure naturally defines a common coordinate system, eliminating the need for post-hoc alignment.

*Stage 1: Per-sample denoising*. Each coupling matrix *U*_*i*_ is eigendecomposed:

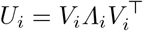

where *V*_*i*_ *∈* ℝ^*F ×F*^ is an orthogonal matrix of eigenvectors and *Λ*_*i*_ = diag(*λ*_*i*,1_, …, *λ*_*i,F*_) with eigenvalues in decreasing order. We retain the top *r*_*i*_ components to form a low-rank approximation:

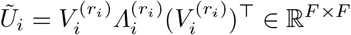

The rank *r*_*i*_ is determined adaptively for each sample using a Kneedle-based elbow detection procedure [54] applied to the log-eigenvalue spectrum. Specifically, let *g*_*j*_ = log(*λ*_*i,j*_) for *j* = 1, …, *F*. We normalize both the index axis *t*_*j*_ = *j* and the log-eigenvalues *g*_*j*_ to the unit interval [0, 1], yielding normalized coordinates 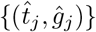. We then compute the perpendicular distance of each point to the line connecting the first and last points 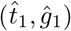 and 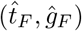:

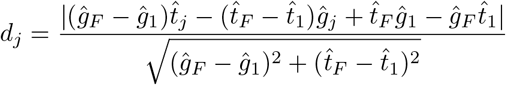

The elbow is defined as *r*_*i*_ = arg max_*j*_ *d*_*j*_, the index at which the log-eigenvalue curve deviates maximally from a linear decay. Although *r*_*i*_ may differ across samples, *Ũ*_*i*_ remains *F × F* for all *i*, so the denoised matrices are directly summable. This step prevents idiosyncratic noise dimensions from accumulating in the subsequent aggregation.

*Stage 2: Aggregation and global basis extraction*. The denoised matrices are summed and eigendecomposed:

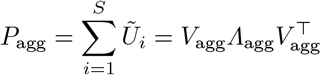

where 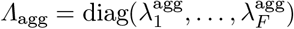 with eigenvalues in decreasing order. We retain the top *d* components to define the shared latent basis:

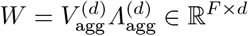

The dimensionality *d* is determined by applying the Kneedle algorithm to 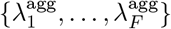. While Stage 1 removes noise within each individual, this global truncation isolates the dominant latent dimensions consistently present across the population.

#### Generation of the Final Feature × Sample Embedding

For each sample *i*, we project its original coupling matrix onto the shared latent basis:

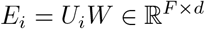

We use the original *U*_*i*_ rather than the denoised *Ũ*_*i*_ because the shared basis *W* already acts as a denoising filter by restricting the projection to consensus dimensions. Applying *W* to *U*_*i*_ thus preserves the full signal content of each individual’s connectivity structure while naturally suppressing sample-specific noise.

The collection of all sample-specific embeddings forms a third-order tensor *ε ∈* ℝ^*F ×d×S*^. The slice 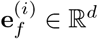 represents the embedding vector of feature *f* in sample *i*, encoding its connectivity profile relative to all other features in the shared latent space. For any feature *f*, the collection 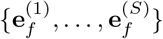 traces how that feature’s network context varies across the cohort, providing the foundation for downstream differential connectivity analysis and sample subgroup detection.

### 4.2 Differential Connectivity Feature Identification

The input consists of the feature × sample embedding tensor *ε∈*ℝ^*F ×d×S*^ produced by the MOSAIC framework (Section 4.1) and a vector of condition labels *c* = (*c*_1_, …, *c*_*S*_) assigning each sample to one of *G* groups.

#### Construction of Feature Trajectory Matrices

For each feature *f ∈ {* 1, …, *F}*, we extract its slice from the tensor to form a feature-specific matrix:

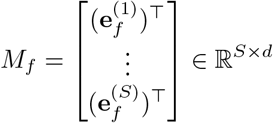

where the *i*-th row 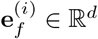 is the embedding of feature *f* in sample *i*. This matrix captures the trajectory of a single feature across all individuals in the cohort: if the feature’s connectivity profile is stable across conditions, the rows corresponding to different groups will overlap in embedding space; if the feature undergoes network rewiring, the rows will separate by condition.

#### Quantification of Geometric Shift via PERMANOVA

To test whether feature *f* exhibits a statistically significant shift in its embedding between conditions, we apply a Permutational Multivariate Analysis of Variance (PERMANOVA) to *M*_*f*_. We first compute a pairwise Euclidean distance matrix *D*_*f*_ *∈* ℝ ^*S×S*^, where 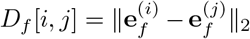. The magnitude of the condition-driven shift is quantified by the pseudo-*F* statistic:

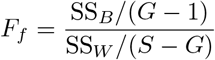

where SS_*B*_ and SS_*W*_ denote the between-group and within-group sums of squared distances, respectively, *S* is the total number of samples, and *G* is the number of condition groups. A large *F*_*f*_ indicates that the embedding positions of feature *f* are more separated between conditions than within conditions, consistent with network rewiring.

#### Statistical Significance via Global Null Model

To distinguish true biological rewiring from artifacts of the embedding procedure, we construct an empirical null distribution using a global permutation strategy. We randomly reassign cell-to-sample labels across the cohort, destroying all condition-specific biological signals while preserving the overall data structure, and recompute the entire MOSAIC pipeline on the permuted data to obtain a null embedding tensor *ε* ^(0)^. We then compute the pseudo-*F* statistic for every feature from this null embedding, yielding a set of null statistics 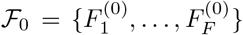. Because no true condition effect exists in the permuted data, these statistics represent the background noise distribution of the full framework. The empirical *p*-value for feature *f* is:

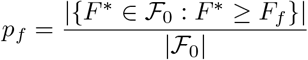

We apply the Benjamini-Hochberg procedure to control the false discovery rate. Features with adjusted *p*-value *<* 0.05 are classified as Differential Connectivity features.

### 4.3 Unsupervised Sample Subgroup Detection

#### Construction of Feature Stratification Profiles

We first characterize the patient-separating behavior of each individual feature. For each feature *f ∈ {*1, …, *F}*, we extract its trajectory matrix *M*_*f*_ *∈* ℝ ^*S×d*^ (as defined in Section 4.2), where the *i*-th row 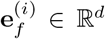 is the embedding of feature *f* in sample *i*. We compute a feature-specific sample-to-sample distance matrix *D*_*f*_ ∈ ℝ^*S×S*^ using cosine distance:

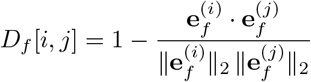

We vectorize the lower triangular portion of *D*_*f*_ into a stratification profile **v**_*f*_ *∈* ℝ^*S*(*S−*1)*/*2^. This vector encodes the complete pairwise patient similarity structure as seen through the lens of feature *f* alone.

#### Identification of Coherent Feature Modules

Features that induce similar patient stratifications are likely to participate in the same underlying biological process. To identify such groups, we construct a feature-feature similarity matrix *Γ*_feat_ *∈*ℝ ^*F ×F*^, where each entry is the Spearman rank correlation between the corresponding stratification profiles:

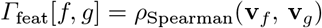

We apply hierarchical clustering to *Γ*_feat_ to identify feature modules *ℳ⊂ {*1, …, *F}*, defined as subsets of features exhibiting highly correlated stratification patterns across the cohort.

#### Module-Driven Patient Clustering

For each identified module *ℳ*, we compute a module-specific patient similarity matrix *P*_*ℳ*_ *∈* ℝ^*S×S*^. The similarity between patients *i* and *j* is defined as the cosine similarity of their aggregated module embeddings:

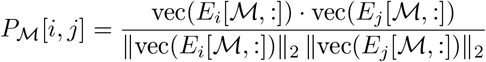

where *E*_*i*_[*ℳ*, :] *∈* ℝ^|*ℳ*|*×d*^ is the submatrix of embeddings for features in *ℳ* for sample *i*, and vec(*·*) denotes vectorization. Hierarchical clustering on *P*_*ℳ*_ reveals patient subgroups driven specifically by the biological process captured within module *ℳ*.

### 4.4 Clinical Outcome Prediction

The feature *×* sample embedding tensor *ε∈*ℝ^*F ×d×S*^ produced by MOSAIC encodes each feature’s connectivity profile per individual. This representation naturally defines a patient-level descriptor suitable for supervised prediction of clinical outcomes.

#### Construction of Patient-Level Connectivity Profiles

For each patient *i*, the embedding slice *E*_*i*_ *∈*ℝ^*F ×d*^ captures the connectivity state of all *F* features in *d* latent dimensions. To convert this matrix into a single feature vector for regression, we vectorize:

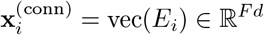

Each entry of 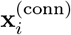 corresponds to one feature–dimension pair, preserving the full topological information of that patient’s regulatory network.

#### Severity Prediction via *L*_1_-Regularized Logistic Regression

Given a binary clinical outcome *y*_*i*_ *∈ {* 0, 1*}* for each patient *i*, we model the log-odds of the positive class as a linear function of the connectivity profile:

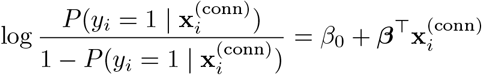

where *β*_0_ *∈*ℝ is the intercept and ***β*** *∈*ℝ^*Fd*^ is the coefficient vector. The parameters are estimated by minimizing the *L*_1_-penalized negative log-likelihood:

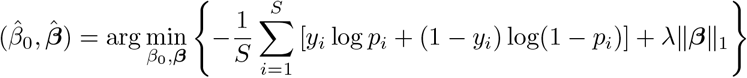

where 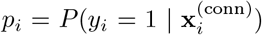 and *λ >* 0 controls the sparsity of the solution. The *L*_1_ penalty induces exact zeros in 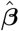, performing simultaneous estimation and feature selection. The predicted risk score for patient *i* is the fitted probability 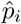.

To quantify the complementarity of connectivity and abundance signals, we additionally train an Abundance model on pseudobulk expression profiles 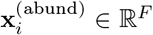 using the same formulation, and an Integrated model via late fusion, where the final risk score is the unweighted mean of the two models’ predicted probabilities.

#### Gene-Level Importance Aggregation

Because each gene *f* contributes *d* entries to the vectorized input, coefficient-based feature selection methods (e.g., Lasso) assign *d* separate coefficients 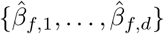 to each gene. To recover gene-level importance, we aggregate across latent dimensions:

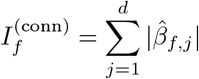

This score quantifies the total contribution of feature *f* ‘s connectivity profile to the prediction, enabling direct comparison with abundance-based importance scores.

### 4.5 Datasets

#### Prefrontal Cortex (PFC) Multi-Omic Cohort

We used a single-nucleus multi-omic dataset of human prefrontal cortex tissue comprising 31 donors (18 HIV+, 13 controls) with paired single-nucleus RNA sequencing (snRNA-seq) and single-nucleus ATAC sequencing (snATAC-seq) profiled from the same nuclei [12]. This cohort is used for validating inter-sample feature relationship variation (Figure 2B), benchmarking cross-sample embedding comparability (Figure 2C–D), and unsupervised subgroup detection (Figure 5). Figure-specific cell type selection, sample filtering, and feature space definitions are described in Sections 4.7 and 4.9.

#### T-Cell CITE-seq Vaccination Cohort

We used a published CITE-seq dataset providing paired measurements of the transcriptome (RNA) and cell-surface proteins (Antibody-Derived Tags, ADT) from human peripheral blood mononuclear cells (PBMCs) of 8 donors, profiled before (Day 0) and after (Day 7) vaccination [13]. This cohort is used for differential connectivity analysis (Figure 4). Cell type filtering and feature selection are described in Section 4.8.

#### COVID-19 scRNA-seq Cohort

We used a large-scale single-cell RNA sequencing dataset of peripheral blood mononuclear cells (PBMCs) from 151 COVID-19 patients [39]. Clinical outcomes were binarized into two severity classes (Moderate and Severe); control samples were excluded. This cohort is used for clinical outcome prediction (Figure 6). Lineage-specific subsetting and feature filtering are described in Section 4.10.

### 4.6 Multi-Modal Simulation Framework

We developed a simulation framework to generate multi-modal single-cell datasets with known groundtruth feature module structure and controlled differential connectivity (DC) signals. The framework serves two purposes: benchmarking feature embedding quality (Figure 2A) and evaluating DC identification sensitivity and robustness (Figure 3). The generative model is shared across all scenarios; what differs is which sources of variation are activated.

#### Generative Model

The simulation generates data for *S* = *S*_*A*_ + *S*_*B*_ samples under conditions A and B across *K* modalities, with *F*_1_, …, *F*_*K*_ features in each modality, respectively. Each sample contains approximately *n* cells with random variation. The latent dimensionality is *r*.

For each sample *i* and modality *k*, the observed data matrix is generated as:

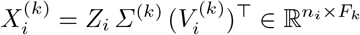

where 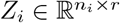 is the cell embedding matrix, *Σ*^(*k*)^ *∈* ℝ^*r×r*^ is a diagonal matrix of singular values, and 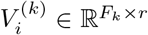 is the feature loading matrix. The three components encode distinct sources of biological variation, described below.

##### Cell embeddings Z_i_

Cells are organized into *C* clusters to mimic cell-type structure. We first draw *C* cluster centers *{* ***µ***_1_, …, ***µ***_*C*_ *} ⊂*ℝ^*r*^ from a standard normal distribution, shared across all samples. For each cell *j* in sample *i*, we assign it to a cluster *c*_*j*_ *∈ {* 1, …, *C}* uniformly at random and generate its embedding as:

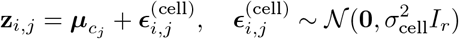

The cell noise level *σ*_cell_ controls within-cluster heterogeneity. Cluster centers are identical across samples, so cell-type composition is shared.

##### Feature loadings 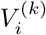

Features are similarly organized into *C* modules using the same set of cluster centers as the cell embeddings, establishing a shared latent geometry between cells and features. For each modality *k*, we first construct a global loading matrix 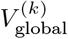 by assigning each feature to a module and generating its loading as the module center plus Gaussian noise with standard deviation *σ*_feature_. Sample-specific loadings are then generated by adding a second layer of noise:

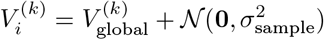

where the noise is applied element-wise. The feature noise *σ*_feature_ controls how tightly features cluster around their module centers (affecting embedding quality), while the sample noise *σ*_sample_ controls inter-individual variation in feature loadings (affecting the degree to which feature relationships differ across samples).

##### Singular values Σ^(k)^

The diagonal matrix *Σ*^(*k*)^ = diag(*σ*_1_, …, *σ*_*r*_) with *r* linearly spaced values is identical across all modalities, ensuring that no modality dominates the latent structure.

#### Differential Connectivity Signal

To simulate differential connectivity, we designate a proportion of features in each modality as DC features. For samples in condition B, the loading rows of DC features are replaced via a constrained permutation: each DC feature’s loading row is swapped with that of a feature belonging to a *different* module. This ensures maximal rewiring: DC features change their network neighborhood between conditions, rather than merely shifting within their original module.

To isolate the connectivity signal from mean expression differences, we apply a mean-correction step: after permutation, the column means of DC features in condition B samples are adjusted to match those of a reference condition A sample. This produces a pure DC signal where features exhibit altered inter-feature relationships between conditions without any change in marginal expression levels.

#### Differential Expression Signal

As a comparator scenario, we also simulate a pure differential expression (DE) signal. In this case, no loading permutation is applied. Instead, the expression values of DE features in condition B samples are multiplied by a fold change *δ*, producing mean shifts without altering feature-feature connectivity structure.

#### Non-Linear Transformation

To test robustness to non-linear distortions that arise in real single-cell data (e.g., from library size normalization, dropout, or count-based noise), we optionally apply a sigmoid transformation to the generated data:

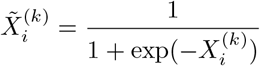

element-wise. This compresses the data into [0, 1] and introduces non-linearity between the latent factors and the observed values, breaking the exact linear SVD structure.

#### Batch Effects

To simulate technical confounders, we optionally add per-feature batch effects. Samples are assigned to *B* batches, balanced across conditions. For each feature *f* in modality *k* and batch *b*, we apply:

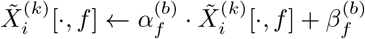

where the multiplicative scale 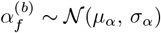 and additive shift 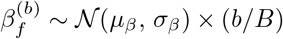 are drawn independently for each feature. The batch-proportional shift ensures that batch effects grow with batch index, mimicking systematic technical drift.

#### Simulation Scenarios

The generative model above is configured differently for each figure:

##### Feature embedding benchmarking

We simulate *S* = 20 samples (10 per condition) across *K* = 2 modalities with *F*_1_ = 1,000 and *F*_2_ = 800 features, approximately 1,000 cells per sample (with random variation of *±* 200), a latent dimensionality of *r* = 10, and *C* = 10 clusters. No DC or DE signal is introduced. The non-linear sigmoid transformation is applied. The singular values are set to *Σ*^(*k*)^ = diag(5, …, 1). The sample-wise and cell-wise noise levels are fixed at *σ*_sample_ = 0.5 and *σ*_cell_ = 0.5, while the feature-wise noise level *σ*_feature_ is progressively increased to create a difficulty gradient for feature module recovery.

##### DC identification benchmarking

We simulate *S* = 20 samples (10 per condition) across *K* = 3 modalities with *F*_1_ = 1,000, *F*_2_ = 800, and *F*_3_ = 600 features, approximately 1,000 cells per sample (with random variation of *±* 200), a latent dimensionality of *r* = 10, and *C* = 10 clusters/modules. 20% of features in each modality are designated as DC features. The singular values are set to *Σ*^(*k*)^ = diag(5, …, 1), and the noise levels are *σ*_sample_ = 0.5, *σ*_cell_ = 0.5, and *σ*_feature_ = 0.5. The non-linear transformation is not applied. Three primary scenarios are evaluated:

**1. Pure Connectivity (DC):** DC features undergo constrained loading permutation with mean correction. This tests whether methods can detect rewiring in the absence of expression changes.

**2. Confounded (DC + DE):** DC features undergo both loading permutation and a multiplicative fold change (*δ* = 2), without mean correction. This tests whether methods can identify connectivity changes when confounded by co-occurring expression shifts.

**3. Mean Shift Only (DE):** Features undergo multiplicative fold change (*δ* = 2) without any loading permutation. This tests specificity: methods should *not* flag these features as DC.

Additional robustness scenarios include: (i) batch effects added to the Pure Connectivity scenario, with *B* = 4 batches (5 samples each), multiplicative scale 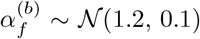, and additive shift 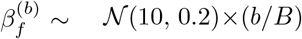, testing whether methods are robust to technical confounders; and (ii) a signal strength sweep, where the singular value matrix *Σ*^(*k*)^ is progressively attenuated from Strong (diag(10, …, 1)) through Medium (diag(5, …, 0.5)) to Weak (diag(0.1, …, 0.01)).

### 4.7 Validation of MOSAIC Feature × Sample Embeddings

We validated MOSAIC’s feature × sample embeddings along three axes: whether MOSAIC produces high-quality feature embeddings (Figure 2A), whether sample-specific embeddings are biologically justified (Figure 2B), and whether MOSAIC’s joint spectral integration produces more comparable cross-sample embeddings than independent embedding followed by post-hoc alignment (Figure 2C–D).

#### Feature Embedding Quality on Simulated Data

Using the multi-modal simulation framework described in Section 4.6 with random seed=42, we generated synthetic multi-modal datasets with known ground-truth feature module assignments. The sample-wise and cell-wise noise levels were fixed at *σ*_sample_ = 0.5 and *σ*_cell_ = 0.5, respectively, while the feature-wise noise level *σ*_feature_ (the standard deviation of additive Gaussian noise applied to feature loadings during data generation) was varied progressively to create a difficulty gradient. At each noise level, we applied four methods to obtain feature embeddings and evaluated their ability to recover the true module structure via *k*-means clustering and the Adjusted Rand Index (ARI). Each noise level was repeated across multiple simulation replicates to obtain stable estimates.

#### Method-specific procedures

For MOSAIC, we ran the full framework (Section 4.1) on the simulated cohort to obtain the sample-specific embedding matrices *{E*_1_, …, *E*_*S*_*}* and averaged across samples to produce a single feature embedding matrix 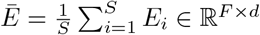, where each feature’s embedding is its corresponding row in *Ē*. For PCA, we pooled cells across all simulated samples, scaled each modality independently to zero mean and unit variance, and concatenated them horizontally into a single cells ×\times × features matrix. We then computed the truncated SVD; the right singular vectors, scaled by their corresponding singular values, served as the feature embedding. For MOFA+, we ran the method in its default multi-group mode on the pooled multi-modal data and extracted the feature weight matrix, where each feature’s loading vector across latent factors constitutes its embedding. For SIMBA, we ran the method with default parameters (50-dimensional embeddings) and extracted the feature embedding vectors from the shared graph embedding space. Each method used its default or recommended dimensionality: MOSAIC determined *d* automatically via the Kneedle algorithm, PCA and MOFA+ use the same Kneedle-based elbow method to select the number of eigenvector or factor to retain, and SIMBA used its default of 50 dimensions.

#### Data Preprocessing and Experimental Setup

The remaining validation experiments use the PFC multi-omic cohort (Section 4.5). For snRNA-seq, data were normalized using log-transformation with a scale factor of 10,000, and the top 10,000 highly variable genes that were also expressed (total count 10) across all retained samples were selected. For snATAC-seq, we applied TF-IDF normalization and retained the top variable features at the 99th percentile quantile. For the cross-sample comparability benchmarking (Figure 2C-D), we selected 6 control donors with the highest cell counts and split each donor’s cells into 10 non-overlapping pseudo-replicates of equal size (random seed = 42). Within each pseudo-replicate set, features were further filtered to retain only those with a total count *>* 3 across all pseudo-replicates; for snRNA-seq, if the number of remaining genes exceeded 3,000, the top 3,000 highly variable genes were selected.

#### Inter-Sample Feature Relationship Variation

For each of the *S* = 31 PFC samples, we computed a feature-feature Pearson correlation matrix *R*_*i*_ *∈* ℝ^*F ×F*^ from the normalized multi-omic expression data, where each entry *R*_*i*_[*j, l*] is the Pearson correlation between features *j* and *l* across the *n*_*i*_ cells in sample *i*. To quantify how much these correlation structures differ across individuals, we computed pairwise Frobenius distances between all 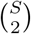 pairs of sample-specific correlation matrices:

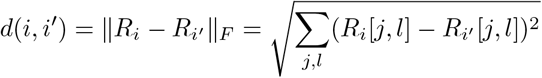

These distances were stratified by modality block (gene–gene, gene–peak, and peak–peak submatrices of *R*_*i*_) to assess whether inter-individual variation is present within and across modalities.

To determine whether the observed variation exceeds what would arise from finite cell sampling alone, we constructed a null distribution by randomly shuffling cell-to-sample assignments across the cohort, destroying all biological differences between individuals while preserving the overall data structure, and recomputing the correlation matrices. We generated 10 null permutations and computed the same pairwise Frobenius distances within each. Significance was assessed by a one-sided Wilcoxon rank-sum test comparing real versus null distances within each modality block.

To illustrate this variation at the level of individual features, we selected nine representative feature pairs (three per modality type: gene–gene, gene–peak, peak–peak) exhibiting high cross-sample variability. Pairs were ranked by the standard deviation of their Pearson correlations across the 31 samples, and the top pairs were selected subject to two constraints: no feature appeared in more than one pair, and peak–peak pairs were restricted to cis-regulatory relationships (features on the same chromosome within 500 kb).

#### Cross-Sample Embedding Comparability Benchmarking

We benchmarked MOSAIC against the same three alternative approaches: PCA, MOFA+, and SIMBA, each representing a different analytical paradigm (baseline dimensionality reduction, multi-modal factor model, and graph-based co-embedding, respectively). Because these methods produce independent per-sample embeddings that reside in separate coordinate systems, each was paired with Generalized Procrustes Analysis (GPA) to align all embeddings into a common coordinate frame before evaluation. GPA avoids the bias of pairwise Procrustes alignment, which depends on an arbitrarily chosen reference configuration. Instead, GPA iteratively estimates a consensus configuration and aligns all matrices to it simultaneously. Let *{Z*_1_, …, *Z*_*N*_*}* denote the *N* = 60 unaligned feature embedding matrices, each in ℝ^*F ×d*^, restricted to the intersection of features shared across all pseudo-replicates. The algorithm proceeds as follows:

1. Initialize the consensus as 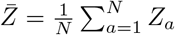.

2. For each *a*, align *Z*_*a*_ to 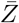 via symmetric Procrustes superimposition, which finds the optimal orthogonal rotation and uniform scaling that minimizes 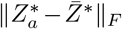, where both *Z*_*a*_ and 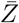 are symmetrically rescaled (denoted by *\**). Replace *Z*_*a*_ with the aligned result.

3. Recompute 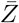 from the aligned matrices. If the change in consensus 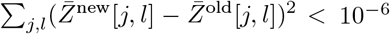, stop; otherwise return to step 2.

Each pairwise alignment was computed using `vegan::procrustes` with `symmetric = TRUE`. MOSAIC produces embeddings that are natively comparable across samples and requires no post-hoc alignment.

##### Pseudo-replicate generation

We selected six control donors from the PFC cohort with the highest cell counts. For each donor, cells were randomly (random seed = 42) partitioned into 10 non-overlapping pseudo-replicates of equal size, yielding 50 pseudo-samples in total. Donor identity served as the ground truth for evaluation, providing a clean validation target that does not conflate methodological quality with biological discovery.

##### Cell count sweep

To assess robustness under varying data sparsity, we repeated the pseudo-replicate construction at multiple cell counts per replicate: *c ∈ {* 200, 300, 400, 500, 600 *}*. At each cell count, the entire pipeline was rerun for all four methods. Each method was run following the same procedure as described above.

##### Evaluation metrics

For each method, we computed pairwise Frobenius distances between the *F × d* embedding matrices of all 60 pseudo-replicates:

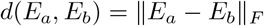

yielding a 60 *×* 60 distance matrix. We evaluated donor identity recovery using the Average Silhouette Score computed on this full-dimensional distance matrix (not on any 2D projection), with donor identity as cluster labels.

### 4.8 T-Cell CITE-seq Differential Connectivity Analysis

#### Data Preprocessing

We utilized the CITE-seq vaccination cohort (Section 4.5), filtering for CD4+ Naive T-cells to isolate the specific signals of T-cell activation. This yielded *S* = 16 samples (8 donors *×* 2 timepoints: Day 0 and Day 7). RNA data were normalized using log-transformation with a scale factor of 10,000; ADT data were normalized using centered log-ratio (CLR) transformation. To ensure embedding stability, we retained only features (genes and proteins) with a total count *>* 3 across all 16 samples.

#### DC Feature Identification

We applied the MOSAIC framework (Section 4.1) to the preprocessed data to generate the feature *×* sample embedding tensor *ε∈*ℝ^*F ×d×S*^, and identified DC features using the PERMANOVA-based procedure (Section 4.2) with condition labels corresponding to Day 0 versus Day 7. Features with Benjamini–Hochberg adjusted *p*-value *<* 0.05 were classified as DC features.

#### Quantification of Network Turnover

To independently validate that DC features undergo greater neighborhood reorganization than background features, we quantified the stability of each feature’s local connectivity across conditions using the Jaccard Index of neighbor overlap. We first computed condition-specific consensus embeddings by averaging across samples within each timepoint:

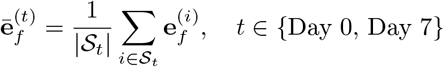

where *𝒮*_*t*_ denotes the set of samples in timepoint *t*. For each feature *f*, we identified its *K* = 20 nearest neighbors in the consensus embedding space for each timepoint, denoted 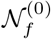 and 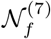, respectively.

The Jaccard Index quantifies the overlap:

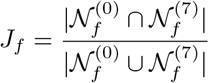

A low *J*_*f*_ indicates substantial rewiring of feature *f* ‘s network neighborhood between conditions. Statistical comparison of Jaccard scores between DC and background features was performed using a Wilcoxon rank-sum test.

#### Validation of Abundance Stability

To confirm that the identified DC features reflect network rewiring rather than abundance changes, we performed a pseudobulk analysis. For each sample *i*, we aggregated single-cell expression counts by computing the mean expression per feature, yielding a pseudobulk vector 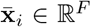. Statistical significance of abundance differences between Day 0 and Day 7 was assessed using a two-sample *t*-test for each feature.

#### Visualization of Embedding Shift

To visualize the geometric shift in a feature’s connectivity profile, we extracted its sample-specific embedding vectors 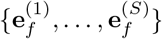 from the tensor *E* and applied Multidimensional Scaling (MDS) to project these *d*-dimensional vectors into a 2D space. Points were colored by timepoint, and condition centroids were connected by an arrow to indicate the direction of the connectivity shift. Network plots (Figure 4E) were generated by visualizing the top *K* = 20 nearest neighbors of the target feature in each condition-specific consensus embedding.

#### Functional Annotation

To characterize the biological coherence of the identified DC features, we performed Reactome pathway enrichment analysis [41] on the full set of 393 DC features. Protein-protein interaction (PPI) network analysis was conducted using the STRING database [55], retaining interactions with a confidence score above the default threshold. The enrichment of observed interactions relative to the expected number was assessed using STRING’s internal hypergeometric test.

### 4.9 PFC Multi-Omic Subgroup Characterization

#### Data Preprocessing

We used the PFC multi-omic cohort (Section 4.5), isolating L2/3 inhibitory neurons for analysis. One sample with fewer than 10 cells in this population was excluded, yielding *S* = 30 donors (18 HIV+, 12 controls). For snRNA-seq, data were normalized using log-transformation with a scale factor of 10,000, and the top 10,000 highly variable genes with a total expression count 3 across all retained samples were selected. For snATAC-seq, we computed gene activity scores using `Signac::GeneActivity` and normalized via log-transformation with a scale factor of 10,000; the top 3,000 variable gene activity features with a total count *>* 5 across all retained samples were selected.

#### Subgroup Validation: HIV+ vs. Control

To validate that MOSAIC’s unsupervised subgroup detection can recover known biological structure, we applied the full framework (Sections 4.1 and 4.3) to all 30 donors, treating disease labels as unknown. The feature module identification and module-driven patient clustering procedures (Section 4.3) were applied to test whether MOSAIC could separate HIV+ from control samples without supervision. To confirm that the identified module captures genuine biological signal rather than technical artifacts, we computed single-cell module activity scores using `Seurat::AddModuleScore` on the original expression data and compared the score distributions between HIV+ and control samples.

#### De Novo Discovery of HIV+ Subgroups

To discover previously unknown subtypes within the HIV+ population, we restricted the MOSAIC pipeline to the 18 HIV+ samples and re-applied the unsupervised subgroup detection workflow (Section 4.3). This analysis identified a feature module that partitioned the HIV+ cohort into two subgroups: HIV-Group1 (*n* = 10) and HIV-Group2 (*n* = 8).

#### Molecular Characterization of HIV+ Subgroups

To characterize the molecular divergence between the two subgroups, we performed differential expression (DE) analysis on the snRNA-seq profiles and differential accessibility (DA) analysis on the snATAC-seq gene activity scores using MAST [32], with subgroup membership as the grouping variable. Features with Benjamini–Hochberg adjusted *p*-value *<* 0.05 and |log_2_ FC |*>* 1 were considered significant. Functional interpretation was performed via Reactome pathway enrichment analysis [41] on the set of genes significantly upregulated in HIV-Group1.

### 4.10 COVID-19 Patient Severity Prediction

#### Data Preprocessing

We used the COVID-19 scRNA-seq cohort (Section 4.5). The dataset was subset into five major immune lineages (Monocytes, B cells, CD4+ T cells, CD8+ T cells, and NK cells) for lineage-specific analysis. For each cell type, patient samples containing fewer than 10 cells were discarded. To ensure a conserved feature space across all individuals, we retained only genes with a total count *>* 3 in every remaining patient sample and kept the intersection of these gene lists. This strict filtering prevents artifacts driven by patient-specific dropout. RNA data were normalized using log-transformation with a scale factor of 10,000 prior to applying MOSAIC.

#### Lineage Screening

To identify the most informative immune compartment, we applied the MOSAIC framework (Section 4.1) independently to each of the five cell types, constructed patient-level connectivity profiles (Section 4.4), and trained Lasso classifiers to predict severity. We compared AUC-ROC across lineages; monocytes achieved the highest predictive accuracy and were selected for all subsequent analyses.

#### Predictive Modeling

Within the monocyte compartment, we trained the Abundance, Connectivity, and Integrated models as formulated in Section 4.4, implemented using the `glmnet` R package (`family = “binomial”`, *α* = 1). Pseudobulk profiles were constructed by aggregating raw UMI counts across all cells per patient, followed by log-transformed counts-per-10,000 (log-CP10K) normalization. For each patient, the risk score is the predicted probability of the Severe class.

Model performance was evaluated using repeated stratified 5-fold cross-validation (repeated 10 times with different random seeds 1:10). Within each training fold, the regularization parameter *λ* was optimized via internal cross-validation to minimize binomial deviance, and predictions on held-out test folds were generated using *λ*_min_. Predictive accuracy was quantified using AUC-ROC, and statistical significance of performance differences between models was assessed using the two-sided DeLong test.

#### Complementary Analysis

To assess the complementarity of abundance- and connectivity-based signals, we compared patient-specific risk scores from the Abundance and Connectivity models. Patients were categorized by their classification pattern relative to the optimal decision threshold of each model: *consensus* (both models agree), *connectivity-rescued* (severe patients correctly identified only by the Connectivity model), *abundance-rescued* (correctly identified only by the Abundance model), and *abundance false positives* (moderate patients misclassified as severe by the Abundance model but correctly classified by the Connectivity model).

#### Molecular Driver Analysis

To identify the molecular features driving each model’s predictions, we extracted non-zero Lasso coefficients from models fitted on the full dataset. Gene-level importance for the Connectivity model was computed using the aggregation procedure in Section 4.4; for the Abundance model, importance was defined as the absolute coefficient. The overlap between the two predictive feature sets was quantified using the Jaccard Index. The top 15 features from each model, ranked by relative importance, were visualized using lollipop plots with functional annotations.

## 4.11 Data Availability

The human CITE-seq vaccination cohort data [13] is available from the Gene Expression Omnibus (GEO) under accession number GSE164378 or via the link: https://atlas.fredhutch.org/nygc/multimodal-pbmc/. The single-nucleus multi-omic prefrontal cortex (PFC) dataset [12] is available from the corresponding authors upon reasonable request. The COVID-19 scRNA-seq cohort data [39] is available from GEO under accession number GSE158055.

## 4.12 Code Availability

The source code for MOSAIC and usage examples are available on GitHub at https://github.com/KlugerLab/MOSAIC.

## 4.13 Acknowledgment

Y.K. is supported by R01DA063148, UM1DA051410, U54AG076043, U54AG079759, U01DA053628, P50CA121974

## A Supplementary Figure

**Figure S1.**
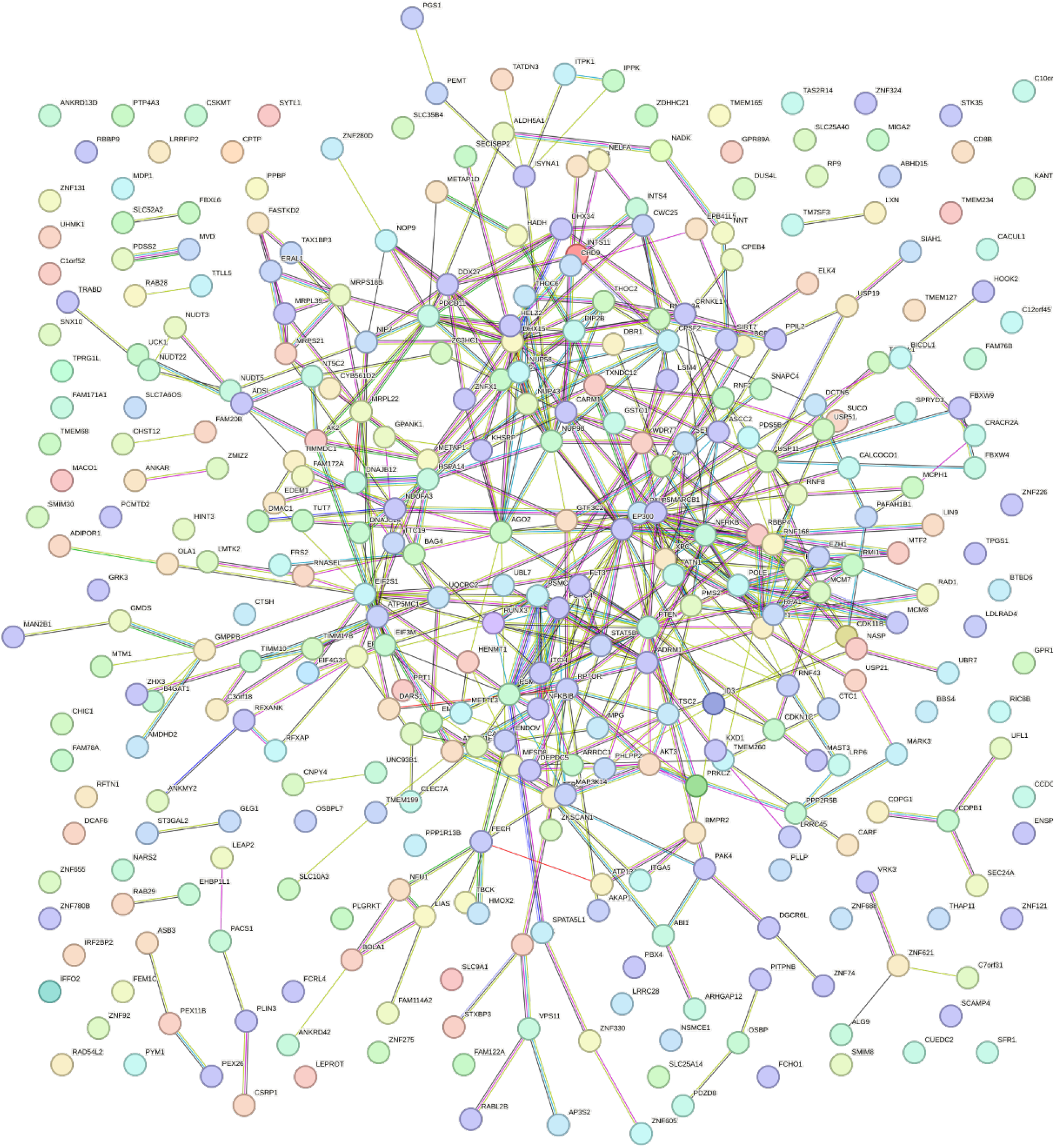
Protein-protein interaction (PPI) network of Differential Connectivity (DC) features. Network visualization of the 393 DC features identified in the T-cell activation dataset, generated using the STRING database. Nodes represent proteins, and colored edges represent known physical or functional interactions. The resulting network exhibits significantly more interactions than expected by chance (493 interactions observed vs. 392 expected; *p* = 5.56 *×* 10^*−*7^). This significant enrichment confirms that the features identified by MOSAIC are not randomly distributed but form a biologically coordinated regulatory network.

**Figure S2.**
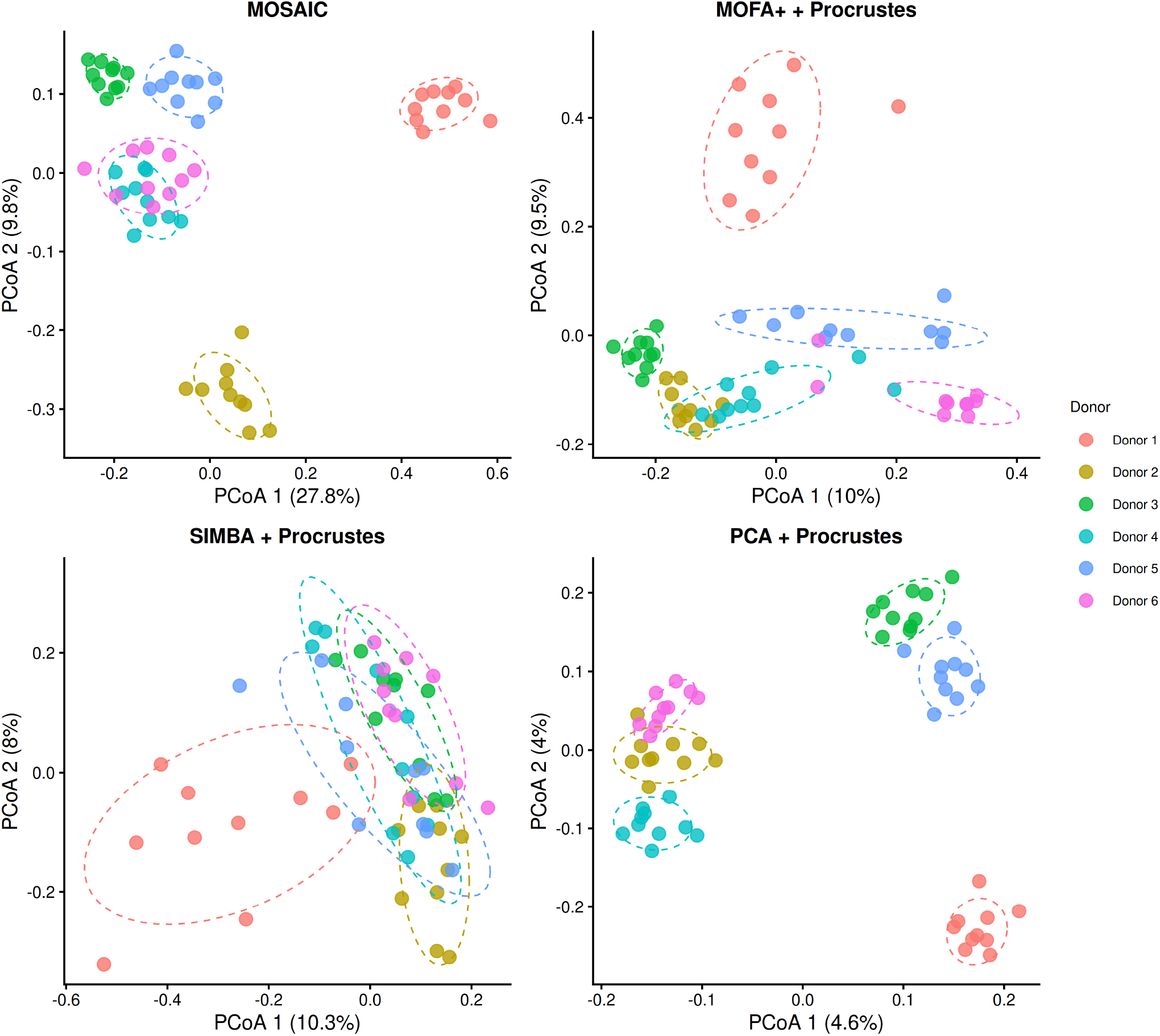
PCoA visualization of cross-sample embedding distances. Principal Coordinates Analysis (PCoA) of pairwise embedding distances at 200 cells per replicate for all four methods: MOSAIC, MOFA+ + Procrustes, SIMBA + Procrustes, and PCA + Procrustes. Points are colored by donor. Axis labels indicate the percentage of variance explained by each coordinate.

## References

1. Fuchou Tang, Catalin Barbacioru, Yangzhou Wang, Ellen Nordman, Clarence Lee, Nanlan Xu, Xiaohui Wang, John Bodeau, Brian B Tuch, Asim Siddiqui, et al. mrna-seq whole-transcriptome analysis of a single cell. Nature methods, 6(5):377–382, 2009.

2. Saiful Islam, Una Kjällquist, Annalena Moliner, Pawel Zajac, Jian-Bing Fan, Peter Lönnerberg, and Sten Linnarsson. Characterization of the single-cell transcriptional landscape by highly multiplex rna-seq. Genome research, 21(7):1160–1167, 2011.

3. Jason D Buenrostro, Paul G Giresi, Lisa C Zaba, Howard Y Chang, and William J Greenleaf. Transposition of native chromatin for fast and sensitive epigenomic profiling of open chromatin, dna-binding proteins and nucleosome position. Nature methods, 10(12):1213–1218, 2013.

4. Ryan M Mulqueen, Dmitry Pokholok, Steven J Norberg, Kristof A Torkenczy, Andrew J Fields, Duanchen Sun, John R Sinnamon, Jay Shendure, Cole Trapnell, Brian J O’Roak, et al. Highly scalable generation of dna methylation profiles in single cells. Nature biotechnology, 36(5):428–431, 2018.

5. Marlon Stoeckius, Christoph Hafemeister, William Stephenson, Brian Houck-Loomis, Pratip K Chattopadhyay, Harold Swerdlow, Rahul Satija, and Peter Smibert. Simultaneous epitope and transcriptome measurement in single cells. Nature methods, 14(9):865–868, 2017.

6. Siddharth S Dey, Lennart Kester, Bastiaan Spanjaard, Magda Bienko, and Alexander Van Oudenaarden. Integrated genome and transcriptome sequencing of the same cell. Nature biotechnology, 33(3):285–289, 2015.

7. Sebastian Pott. Simultaneous measurement of chromatin accessibility, dna methylation, and nucleosome phasing in single cells. eLife, 6:e23203, jun 2017.

8. Richard K Perez, M Grace Gordon, Meena Subramaniam, Min Cheol Kim, George C Hartoularos, Sasha Targ, Yang Sun, Anton Ogorodnikov, Raymund Bueno, Andrew Lu, et al. Single-cell rna-seq reveals cell type–specific molecular and genetic associations to lupus. Science, 376(6589):eabf1970, 2022.

9. Hansruedi Mathys, Jose Davila-Velderrain, Zhuyu Peng, Fan Gao, Shahin Mohammadi, Jennie Z Young, Madhvi Menon, Liang He, Fatema Abdurrob, Xueqiao Jiang, et al. Single-cell transcriptomic analysis of alzheimer’s disease. Nature, 570(7761):332–337, 2019.

10. Parker C Wilson, Yoshiharu Muto, Haojia Wu, Anil Karihaloo, Sushrut S Waikar, and Benjamin D Humphreys. Multimodal single cell sequencing implicates chromatin accessibility and genetic background in diabetic kidney disease progression. Nature communications, 13(1):5253, 2022.

11. Julia Gamache, Daniel Gingerich, E Keats Shwab, Julio Barrera, Melanie E Garrett, Cordelia Hume, Gregory E Crawford, Allison E Ashley-Koch, and Ornit Chiba-Falek. Integrative single-nucleus multi-omics analysis prioritizes candidate cis and trans regulatory networks and their target genes in alzheimer’s disease brains. Cell & Bioscience, 13(1):185, 2023.

12. Junchen Yang, Kriti Agrawal, Jay Stanley III, Ruiqi Li, Nicholas Jacobs, Haowei Wang, Chang Lu, Rihao Qu, Declan Clarke, Yuhang Chen, et al. Multi-omic characterization of hiv effects at single cell level across human brain regions. bioRxiv, 2025.

13. Yuhan Hao, Stephanie Hao, Erica Andersen-Nissen, William M Mauck, Shiwei Zheng, Andrew Butler, Maddie J Lee, Aaron J Wilk, Charlotte Darby, Michael Zager, et al. Integrated analysis of multimodal single-cell data. Cell, 184(13):3573–3587, 2021.

14. Adam Gayoso, Romain Lopez, Zoë Steier, Jeffrey Regier, Aaron Streets, and Nir Yosef. A joint model of rna expression and surface protein abundance in single cells. biorxiv, page 791947, 2019.

15. Joshua D Welch, Velina Kozareva, Ashley Ferreira, Charles Vanderburg, Carly Martin, and Evan Z Macosko. Single-cell multi-omic integration compares and contrasts features of brain cell identity. Cell, 177(7):1873–1887, 2019.

16. Manqi Zhou, Hao Zhang, Zilong Bai, Dylan Mann-Krzisnik, Fei Wang, and Yue Li. Single-cell multi-omics topic embedding reveals cell-type-specific and covid-19 severity-related immune signatures. Cell Reports Methods, 3(8), 2023.

17. Yang Xu, Priyojit Das, and Rachel Patton McCord. Smile: mutual information learning for integration of single-cell omics data. Bioinformatics, 38(2):476–486, 2022.

18. Tal Ashuach, Mariano I Gabitto, Rohan V Koodli, Giuseppe-Antonio Saldi, Michael I Jordan, and Nir Yosef. Multivi: deep generative model for the integration of multimodal data. Nature Methods, 20(8):1222–1231, 2023.

19. Boying Gong, Yun Zhou, and Elizabeth Purdom. Cobolt: integrative analysis of multimodal single-cell sequencing data. Genome biology, 22(1):351, 2021.

20. Yingxin Cao, Laiyi Fu, Jie Wu, Qinke Peng, Qing Nie, Jing Zhang, and Xiaohui Xie. Sailer: scalable and accurate invariant representation learning for single-cell atac-seq processing and integration. Bioinformatics, 37(Supplement_1):i317–i326, 2021.

21. Lei Xiong, Kang Tian, Yuzhe Li, Weixi Ning, Xin Gao, and Qiangfeng Cliff Zhang. Online single-cell data integration through projecting heterogeneous datasets into a common cell-embedding space. Nature Communications, 13(1):6118, 2022.

22. Qiuyue Yuan and Zhana Duren. Inferring gene regulatory networks from single-cell multiome data using atlas-scale external data. Nature Biotechnology, 43(2):247–257, 2025.

23. Carmen Bravo González-Blas, Seppe De Winter, Gert Hulselmans, Nikolai Hecker, Irina Matetovici, Valerie Christiaens, Suresh Poovathingal, Jasper Wouters, Sara Aibar, and Stein Aerts. Scenic+: single-cell multiomic inference of enhancers and gene regulatory networks. Nature methods, 20(9):1355–1367, 2023.

24. Junyao Jiang, Pin Lyu, Jinlian Li, Sunan Huang, Jiawang Tao, Seth Blackshaw, Jiang Qian, and Jie Wang. Irena: Integrated regulatory network analysis of single-cell transcriptomes and chromatin accessibility profiles. Iscience, 25(11), 2022.

25. Lihua Zhang, Jing Zhang, and Qing Nie. Direct-net: an efficient method to discover cis-regulatory elements and construct regulatory networks from single-cell multiomics data. Science advances, 8(22):eabl7393, 2022.

26. Huidong Chen, Jayoung Ryu, Michael E Vinyard, Adam Lerer, and Luca Pinello. Simba: single-cell embedding along with features. Nature Methods, 21(6):1003–1013, 2024.

27. Zhi-Jie Cao and Ge Gao. Multi-omics single-cell data integration and regulatory inference with graph-linked embedding. Nature Biotechnology, 40(10):1458–1466, 2022.

28. Ricard Argelaguet, Damien Arnol, Danila Bredikhin, Yonatan Deloro, Britta Velten, John C Marioni, and Oliver Stegle. Mofa+: a statistical framework for comprehensive integration of multi-modal single-cell data. Genome biology, 21(1):111, 2020.

29. Ziqi Zhang, Chengkai Yang, and Xiuwei Zhang. scdart: integrating unmatched scrna-seq and scatac-seq data and learning cross-modality relationship simultaneously. Genome biology, 23(1):139, 2022.

30. Michael I Love, Wolfgang Huber, and Simon Anders. Moderated estimation of fold change and dispersion for rna-seq data with deseq2. Genome biology, 15(12):550, 2014.

31. Mark D Robinson, Davis J McCarthy, and Gordon K Smyth. edger: a bioconductor package for differential expression analysis of digital gene expression data. bioinformatics, 26(1):139–140, 2010.

32. Greg Finak, Andrew McDavid, Masanao Yajima, Jingyuan Deng, Vivian Gersuk, Alex K Shalek, Chloe K Slichter, Hannah W Miller, M Juliana McElrath, Martin Prlic, et al. Mast: a flexible statistical framework for assessing transcriptional changes and characterizing heterogeneity in single-cell rna sequencing data. Genome biology, 16(1):278, 2015.

33. Matthew E Ritchie, Belinda Phipson, D. Wu, Yifang Hu, Charity W Law, Wei Shi, and Gordon K Smyth. limma powers differential expression analyses for rna-sequencing and microarray studies. Nucleic acids research, 43(7):e47–e47, 2015.

34. Chen Weng, Anniya Gu, Shanshan Zhang, Leina Lu, Luxin Ke, Peidong Gao, Xiaoxiao Liu, Yuntong Wang, Peinan Hu, Dylan Plummer, et al. Single cell multiomic analysis reveals diabetes-associated β-cell heterogeneity driven by hnf1a. Nature communications, 14(1):5400, 2023.

35. William S Chen, Nevena Zivanovic, David Van Dijk, Guy Wolf, Bernd Bodenmiller, and Smita Krishnaswamy. Uncovering axes of variation among single-cell cancer specimens. Nature methods, 17(3):302–310, 2020.

36. Mehdi Joodaki, Mina Shaigan, Victor Parra, Roman D Bülow, Christoph Kuppe, David L Hölscher, Mingbo Cheng, James S Nagai, Michaël Goedertier, Nassim Bouteldja, et al. Detection of patient-level distances from single cell genomics and pathomics data with optimal transport (pilot). Molecular systems biology, 20(2):57–74, 2024.

37. Zexuan Wang, Qipeng Zhan, Shu Yang, Shizhuo Mu, Jiong Chen, Sumita Garai, Patryk Orzechowski, Joost Wagenaar, and Li Shen. Qot: efficient computation of sample level distance matrix from single-cell omics data through quantized optimal transport. bioRxiv, 2024.

38. Haidong Yi and Natalie Stanley. sclkme: A landmark-based approach for generating multi-cellular sample embeddings from single-cell data. bioRxiv, 2023.

39. Xianwen Ren, Wen Wen, Xiaoying Fan, Wenhong Hou, Bin Su, Pengfei Cai, Jiesheng Li, Yang Liu, Fei Tang, Fan Zhang, et al. Covid-19 immune features revealed by a large-scale single-cell transcriptome atlas. Cell, 184(7):1895–1913, 2021.

40. Yuhan Hao, Tim Stuart, Madeline H Kowalski, Saket Choudhary, Paul Hoffman, Austin Hartman, Avi Srivastava, Gesmira Molla, Shaista Madad, Carlos Fernandez-Granda, et al. Dictionary learning for integrative, multimodal and scalable single-cell analysis. Nature biotechnology, 42(2):293–304, 2024.

41. Guangchuang Yu and Qing-Yu He. Reactomepa: an r/bioconductor package for reactome pathway analysis and visualization. Molecular BioSystems, 12(2):477–479, 2016.

42. Reinhold Förster, Andreas Schubel, Dagmar Breitfeld, Elisabeth Kremmer, Ingrid Renner-Müller, Eckhard Wolf, and Martin Lipp. Ccr7 coordinates the primary immune response by establishing functional microenvironments in secondary lymphoid organs. Cell, 99(1):23–33, 1999.

43. Fabio Malavasi, Silvia Deaglio, Enza Ferrero, Ada Funaro, Jaime Sancho, Clara M Ausiello, Erika Ortolan, Tiziana Vaisitti, Mercedes Zubiaur, Giorgio Fedele, et al. Cd38 and cd157 as receptors of the immune system: a bridge between innate and adaptive immunity. Molecular medicine, 12(11-12):334–341, 2006.

44. Ann Katrin Greifenberg, Dana Hönig, Kveta Pilarova, Robert Düster, Koen Bartholomeeusen, Christian A Bösken, Kanchan Anand, Dalibor Blazek, and Matthias Geyer. Structural and functional analysis of the cdk13/cyclin k complex. Cell reports, 14(2):320–331, 2016.

45. Kaoru Sugasawa, Jessica MY Ng, Chikahide Masutani, Shigenori Iwai, Peter J van der Spek, André PM Eker, Fumio Hanaoka, Dirk Bootsma, and Jan HJ Hoeijmakers. Xeroderma pigmentosum group c protein complex is the initiator of global genome nucleotide excision repair. Molecular cell, 2(2):223–232, 1998.

46. Yulia V Surovtseva, Dmitri Churikov, Kara A Boltz, Xiangyu Song, Jonathan C Lamb, Ross Warrington, Katherine Leehy, Michelle Heacock, Carolyn M Price, and Dorothy E Shippen. Conserved telomere maintenance component 1 interacts with stn1 and maintains chromosome ends in higher eukaryotes. Molecular cell, 36(2):207–218, 2009.

47. Jian-Xin Lin, Meili Ge, Cheng-yu Liu, Ronald Holewinski, Thorkell Andresson, Zu-Xi Yu, Tesfay Gebregiorgis, Rosanne Spolski, Peng Li, and Warren J Leonard. Tyrosine phosphorylation of both stat5a and stat5b is necessary for maximal il-2 signaling and t cell proliferation. Nature Communications, 15(1):7372, 2024.

48. Anthony Tubbs and André Nussenzweig. Endogenous dna damage as a source of genomic instability in cancer. Cell, 168(4):644–656, 2017.

49. Erica A Mendes, Yuyang Tang, and Guochun Jiang. The integrated stress response signaling during the persistent hiv infection. Iscience, 26(12), 2023.

50. C Akay, KA Lindl, N Shyam, B Nabet, Y Goenaga-Vazquez, J Ruzbarsky, Y Wang, DL Kolson, and KL Jordan-Sciutto. Activation status of integrated stress response pathways in neurones and astrocytes of hiv-associated neurocognitive disorders (hand) cortex. Neuropathology and applied neurobiology, 38(2):175–200, 2012.

51. Charles A Hoeffer and Eric Klann. mtor signaling: at the crossroads of plasticity, memory and disease. Trends in neurosciences, 33(2):67–75, 2010.

52. Anke Di, Shiqin Xiong, Zhiming Ye, RK Subbarao Malireddi, Satoshi Kometani, Ming Zhong, Manish Mittal, Zhigang Hong, Thirumala-Devi Kanneganti, Jalees Rehman, et al. The twik2 potassium efflux channel in macrophages mediates nlrp3 inflammasome-induced inflammation. Immunity, 49(1):56–65, 2018.

53. Qiuyue Yuan and Zhana Duren. Integration of single-cell multi-omics data by regression analysis on unpaired observations. Genome Biology, 23(1):160, 2022.

54. Ville Satopaa, Jeannie Albrecht, David Irwin, and Barath Raghavan. Finding a” kneedle” in a haystack: Detecting knee points in system behavior. In 2011 31st international conference on distributed computing systems workshops, pages 166–171. IEEE, 2011.

55. Christian von Mering, Martijn Huynen, Daniel Jaeggi, Steffen Schmidt, Peer Bork, and Berend Snel. String: a database of predicted functional associations between proteins. Nucleic acids research, 31(1):258–261, 2003.

